# Towards 3D-Bioprinting of an Endocrine Pancreas: A Building-Block Concept for Bioartificial Insulin-Secreting Tissue

**DOI:** 10.1101/2021.02.27.433164

**Authors:** Gabriel Alexander Salg, Eric Poisel, Matthias Neulinger Munoz, Daniel Cebulla, Vitor Vieira, Catrin Bludszuweit-Philipp, Felix Nickel, Ingrid Herr, Nathalia A. Giese, Thilo Hackert, Hannes Goetz Kenngott

**Affiliations:** University Hospital Heidelberg, Department of General, Visceral and Transplantation Surgery, Im Neuenheimer Feld 420, 69120 Heidelberg, Germany; ASD Advanced Simulation and Design GmbH, Erich-Schlesinger-Strasse 50, 18059 Rostock, Germany; INOVA DE GmbH, Im Neuenheimer Feld 515, 69120 Heidelberg, Germany

**Keywords:** bioprinting, tissue engineering, endocrine pancreas, next-generation sequencing, diabetes

## Abstract

**Background & Aims:** 3D-Bioprinting of an endocrine pancreas is a promising future curative treatment for selected patients with insulin secretion deficiency. In this study we present an end-to-end integrative, scalable concept extending from the molecular to the macroscopic level.

**Methods:** A hybrid scaffold device was manufactured by 3D-(bio)printing. INS-1 cells with/without endothelial cells were bioprinted in gelatin methacrylate blend hydrogel. Polycaprolactone was 3D-printed and heparin-functionalized as structural scaffold component. In vitro evaluation was performed by viability and growth assays, total mRNA sequencing, and glucose-stimulated insulin secretion. In vivo, xenotransplantation to fertilized chicken eggs was used to investigate vascularization and function, and finite element analysis modeling served to detect boundary conditions and applicability for human islets of Langerhans.

**Results:** Insulin-secreting pseudoislets were formed and resulted in a viable and proliferative experimental model. Transcriptomics revealed upregulation of proliferative and β-cell-specific signaling cascades, downregulation of apoptotic pathways, and overexpression of extracellular matrix proteins and VEGF induced by pseudoislet formation and 3D culture. Co-culture with human endothelial cells created a natural cellular niche resulting in enhanced insulin response after glucose stimulation. Survival and function of the pseudoislets after explantation and extensive scaffold vascularization of both the hydrogel and heparinized polycaprolactone components were demonstrated *in ovo.* Computer simulations of oxygen, glucose, and insulin flows were used to evaluate scaffold architectures and Langerhans islets at a future transplantation site along neurovascular structures.

**Conclusion:** A defined end-to-end process for multidisciplinary bioconvergence research on a bioartificial endocrine pancreas was developed. A modular, patient-specific device architecture is proposed for future research studies.

## Introduction

Transplantation of islets of Langerhans to selected patients with type 1 and type 3c diabetes mellitus (DM) is an established treatment option.^1, 2^ Autologous transplantation can be performed after isolation of islets from the resected pancreas without the need for life-long immunosuppression, whereas islet transplants for patients with type 1 DM rely on allogenic donor islets.^2^ Current therapeutic limitations include a shortage of donor material but also a substantial loss of islets and impaired long-term function post transplantation.^1, 3-5^ Scaffold-based tissue engineering approaches extend the range of possible transplantation sites and might present a long-term curative treatment.^3, 4, 6-9^ For a successful translational approach, several requirements and properties of a functional tissue-engineered device have to be considered. The scaffold material itself should not induce cytotoxicity or extensive foreign body response and should preferably support or promote rapid vascularization.^10, 11^ Furthermore, the scaffold should be retrievable and possess a certain mechanical strength, at least until tissue remodeling has occured.^10, 11^

Previously, we found that scaffold-based tissue engineering is still hampered by reduced vascularization, causing insufficient nutrition, hypoxia, and immunological host-graft reactions.^3^ The multitude of studies focusing mostly on aspects of the tissue engineering network have not yet provided structured evidence to define a gold-standard approach. Investigations of a variety of different cells, scaffold materials, fabrication techniques, and transplantation sites have not yet consolidated into an entire process leading towards bioartificial organs. In an integrated, multilevel approach, tissue-engineered building blocks are investigated on a molecular level against the background of a macroscopic device for translation to further steps. The end-to-end concept presented here aims to address the challenges of hybrid scaffold fabrication, cellular integration, and functional evaluation to provide experimental proof of function for a 3D-bioprinted hybrid scaffold for insulin-secreting cells that is designed for clinical application.

## Materials and Methods

A detailed account of the materials and methods can be found in the Supplementary Materials.

### Computer-aided design (CAD) model creation and slicing for hybrid scaffold fabrication

CAD models were converted to the G programming language with the Cura software package (v4.1, Ultimaker, Utrecht, Netherlands) and used for polycaprolactone (PCL) outer shell 3D-printing. An integrated slicing software was applied (CellInk, Gothenburg, Sweden) for hydrogel 3D-bioprinting.

### 3D-Printing of PCL, heparin surface functionalization, growth factor addition for hybrid scaffold

PCL components (Facilan™ PCL, 3D4Makers, Haarlem, Netherlands) were fabricated using a dual-extrusion-based 3D-printer (UM S5, Ultimaker) and sacrificial support material. Heparin was covalently bonded to the PCL surface by means of carbodiimide chemistry. Basic fibroblast growth factor (bFGF) or nerve growth factor (NGF) bound to heparinized PCL by immersion. PCL scaffolds, heparin-coated PCL scaffolds and heparin-coated PCL scaffolds with bFGF/NGF addition were analyzed by scanning electron microscopy (Zeiss Leo Gemini 1530, Carl Zeiss, Oberkochen, Germany).

### Cell culture

The rat INS-1 832/3 cell line (insulinoma cell line stably transfected with human insulin; hereinafter INS-1) was obtained from Merck. A HUVEC (human umbilical vein endothelial cell) line was obtained from the American Type Culture Collection (Manassas, USA). INS-1 cells were cultivated in RPMI-1640 supplemented with 10% fetal bovine serum, 1% Geneticin, 1% HEPES 1 M, 1% sodium pyruvate 100 mM, and 0.1% 2-mercaptoethanol. HUVEC were cultivated in Endothelial Cell Growth Medium (Cell Applications, San Diego, USA) supplemented with 1% penicillin/streptomycin and 5% fetal bovine serum.

### Bioprinting of cell-laden hydrogels for hybrid scaffold

Bioprinting was performed using the BioX (CellInk). The cells, either INS-1 only or INS-1 with HUVEC in 1:2 ratio, were gently mixed 1:10 with GelXA LAMININK-411 hydrogel (Lot# IK-3X2123, CellInk). Ultraviolet (UV) crosslinking of the bioprinted structures was performed at 405 nm at 5 cm distance.

### Detection of metabolic activity and proliferation

For a visual assessment of metabolic activity, INS-1 cells in bioprinted grid scaffolds were stained with thiazolyl blue tetrazolium bromide (MTT) after 5 days in culture according to the manufacturer’s protocol (Merck, Darmstadt, Germany). Proliferation was determined using an ATP-based assay with luminescent readout (CellTiter-Glo^®^ 3D Cell Viability Assay, Promega, Walldorf, Germany) according to the manufacturer’s protocol.

### RNA sequencing

Genome-wide expression profiling was provided by the European Molecular Biology Laboratory (EMBL, Heidelberg, Germany). After 5 days in culture, total RNA was isolated from 2Dmonolayer culture and 3D hydrogel culture group (biological replicates) using a RNeasy Mini kit (Qiagen, Hilden, Germany) according to the manufacturer’s instructions. Further information on sample preparation, quality control, library preparation, data curation, and analysis can be found in the Supplemental Materials. Using the DESeq2 and log2 fold change pre-ranked differentially expressed genes, a gene set enrichment analysis was performed using the hallmark gene sets. Additional data analysis was performed using Ingenuity Pathway Analysis (IPA; Ingenuity Systems, Qiagen). Canonical pathway analysis identified the pathways referenced in the Ingenuity Knowledge Base of canonical pathways (11/2020) that were significantly *(P* < .05) altered in the dataset.

### Xenotransplantation to the chorioallantoic membrane (CAM) of fertilized chicken eggs

As described before,^12^ fertilized eggs were obtained from a local ecological hatchery (Gefluegelzucht Hockenberger, Eppingen, Germany). On day 9 of embryonic development, PCL scaffold groups or bioprinted xenografts were placed on the CAM. For explantation, the chicks were ethically euthanized at day 18 of development, 3 days before hatching. PCL scaffolds and bioprinted xenografts were excised. Images of PCL scaffold groups were analyzed using the WimCAM software (Wimasis, Onimagin Technologies, Cordoba, Spain).

### Immunohistochemistry of xenograft tissue

Bioprinted xenografts were fixed, transferred to ethanol, and paraffin embedded (see Supplementary Materials). Whole-slide scans of hematoxylin-eosin (H&E), anti-insulin, and anti-chicken-CD34 stainings were performed (see Supplementary Materials) to identify vascular structures and analyze pseudoislet survival and function in xenografts. Islets were segmented with ilastik using supervised machine learning.

### Glucose-stimulated insulin secretion (GSIS)

INS-1 cells were stained with red fluorescent membrane-inserting dye PKH-26 (Lot# SLBW0232, Merck, Darmstadt, Germany) according to the manufacturer’s protocol. The insulin secretion of 3D-bioprinted INS-1 and INS-1/HUVEC co-culture groups, INS-1 cells seeded on PCL/heparin-PCL scaffolds, and a 2D-monolayer control group was measured. GSIS was initiated by rinsing the cells once with basal-concentration-glucose solution (1.67 mM D-glucose), followed by incubation for 1 h in 1 mL basal-glucose solution. After that, either 1 mL of basal-glucose solution or 1 mL of high-glucose solution (16.7 mM D-glucose) was added to the well, followed by incubation for 2 h. Insulin concentration was determined by chemiluminescence immunoassay (ADVIA CENTAUR, Siemens, Malvern, USA). The total amount of insulin measured for the 3D-bioprinted INS-1 and INS-1/HUVEC co-culture groups and the 2D-monolayer control group was normalized to the respective cell number. INS-1 cells seeded on PCL/heparin-PCL scaffolds and the 2D-monolayer control group were normalized to total protein, and a conversion factor between cell number and total protein was calculated to enable inter-group comparability.

### Computer-aided applicability screening of scaffold architecture by finite element analysis

Diffusion of oxygen, glucose, and secreted insulin through islets of Langerhans encapsuled in a hydrogel shell was modeled using a custom python script (v3.8, https://www.python.org) for input parameterbased insulin secretion based on literature data^13^. The simulations were performed in 2D. Results were extrapolated to 3D spherical setups. In the simulation, diffusion started from outside the capsula and triggered consumption of glucose and oxygen within the islets. Based on simulation results, cell viability was evaluated by considering a minimum local oxygen partial pressure of 0.07 mmHg as necessary for cell survival.

### Statistical analysis

Data analysis and statistical testing was done using R version 3.6.1 and the ggplot2 package. Proliferation assay, vascular ingrowth assay, and GSIS data were analyzed using the nonparametric Wilcoxon rank-sum test. Conditions in 2D and 3D GSIS were normalized to cell number, and PCL scaffold conditions were normalized to total protein. Values not within the *2s* interval were seen as outliers and removed prior to analysis. The results are presented as standard error of the mean (SEM). Sequencing was performed using two biological replicates; other experiments were repeated at least three times. Using IPA, the activity, enrichment and statistical significance of canonical pathways in the dataset was calculated in two ways: (1) ratio of dataset molecule number mapped to pathway divided by total molecule number mapped to canonical pathway; and (2) right-tailed Fisher’s exact test to calculate the probability that the association between dataset genes and canonical pathway is explained by chance alone. Dataset molecules meeting the log fold change cut-off of < −0.5 or > 0.5 and p-value < 0.05 were considered for the analysis.

Statistical significance is depicted with asterisks. The p-values are given as \**P* < .05, \*\**P* < .01, *\*\*\*P* < .001.

## Results

The experimental concept of this study was subdivided into hybrid scaffold fabrication, cellular integration, and functional evaluation of the macrodevice. 3D-Printing was used for fabrication of an outer shell PCL component. The solid polymer component was surface-heparinized, leading to enhanced cell adhesion and hydrophilicity. 3D-Bioprinting was used to manufacture the inner, cell-encapsulating hydrogel structure of the hybrid scaffold. Cellular integration of INS-1 cells in the hydrogel led to formation of proliferative pseudoislets. RNA sequencing revealed upregulation of proliferative and β-cell-specific signaling, downregulation of apoptotic marker genes, but also transient metabolic stress post printing. Functional evaluation of the building blocks was performed by GSIS and analysis of viability, function, and vascularization of xenografts *in ovo.* Hereby, the creation of a natural cellular niche by co-culture of INS-1 with endothelial cells (EC) resulted in enhanced insulin secretion upon glucose stimulation. Functional evaluation of the concept was performed by computer simulation of Langerhans islets.

### 3D-Printing and heparin functionalization of PCL improved biocompatibility

The FDA-approved PCL was chosen as the solid polymer component, the outer shell, of the building blocks, based on criteria such as biocompatibility, mechanical properties, retrievability, potential for modification and functionalization with supplementary substrates, and fabrication of a suitable scaffold architecture within a scalable process.^11^ 3D-Printing with additional sacrificial support material (polyvinyl alcohol) enabled fabrication of precise, complex geometries (Fig. 1A, Appendix S1A). Due to hydrophobic characteristics, further modification by heparin conjugation was performed prior to cell seeding to facilitate cell adhesion and proliferation. Covalent binding of heparin (Hep-PCL group) is shown by scanning electron microscopy compared with untreated controls (Fig. 1B-D).

**Figure 1:**
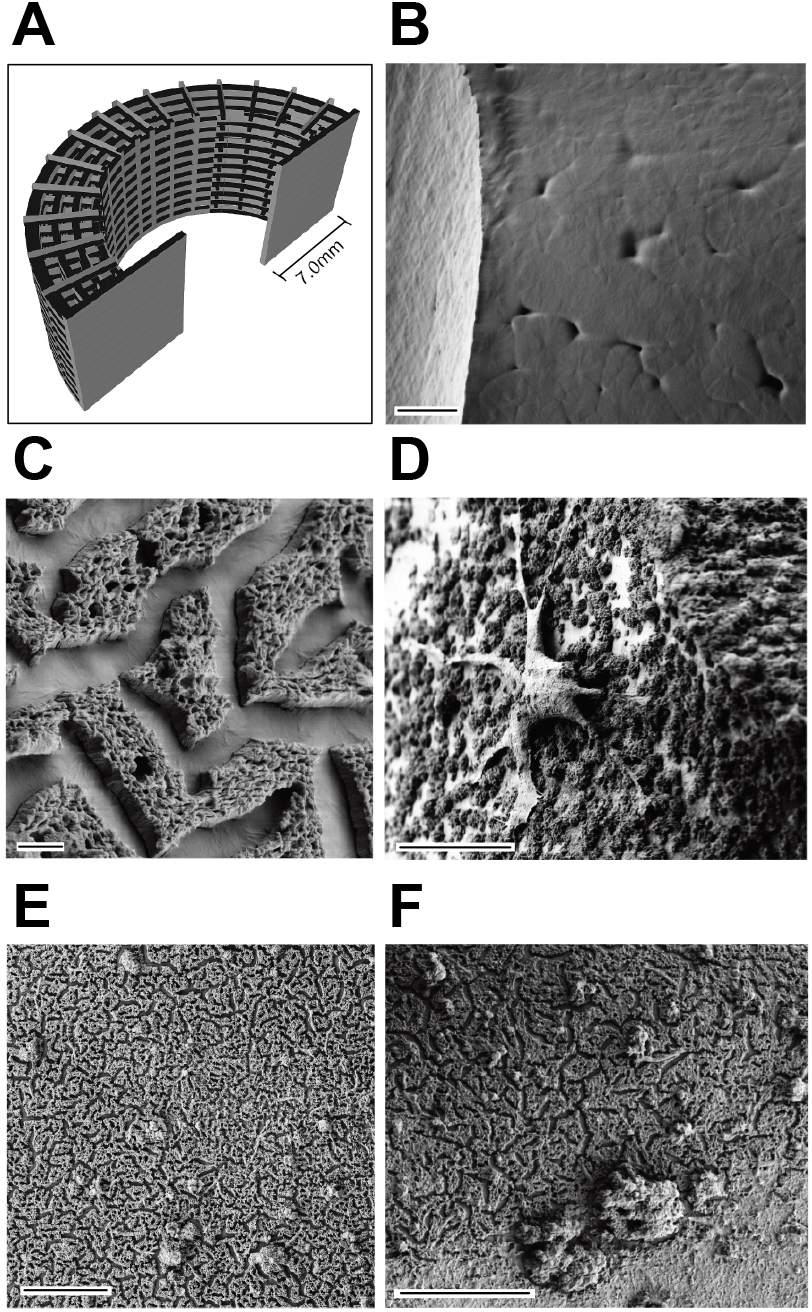
Solid polymer scaffold component was functionalized to enhance biocompatibility, cell adhesion and vascular ingrowth. (A) CAD model of PCL scaffold for 3D-printing as used in the experiments. (B) Scanning electron microscopy of untreated PCL scaffold structure after 3D-printing shows smooth surface with minimal porosity. (C) Surface modification of 3D-printed PCL scaffold with heparin. (D) Adhesion of EC on 3D-printed, heparinized PCL scaffold. (E) Attachment of bFGF and (F) attachment of NGF on 3D-printed, heparinized PCL scaffold validates capability for individualization of the scaffold. Scale bar (B, D-F) 20 μm, (C) 2 μm.

Functionalization remained stable over time (4 weeks, storage in PBS; not displayed). The possibility of further modification of building block components by addition of growth factors is shown using bFGF and NGF (Fig. 1E-F). Both of these growth factors have previously been reported to have beneficial effects on the viability and function of insulin-secreting cells in tissue engineering applications.^8, 9^ We investigated the ability of 3D-printed PCL to support cell adhesion and function in 2D cell culture using INS-1 and HUVEC. INS-1 adhesion could be observed on untreated and heparinized scaffolds, whereas HUVEC adhesion was observed only in the Hep-PCL group (Fig. 1D). In addition to qualitative investigation for biocompatibility of the (Hep-)PCL scaffold, functional analysis showed that INS-1 cells cultured on 3D-printed polymer remained glucose-responsive in all experimental groups. Interestingly, increased insulin secretion after stimulation with glucose was observed in the Hep-PCL group (Appendix S1B).

### Bioprinting of insulin-secreting cells leads to pseudoislet morphology, viability, and growth of bioprinted insulin-secreting cells

In addition to a solid polymer component, the inner core of each building block contains a cell-encapsulating hydrogel. 3D-Bioprinting of INS-1 cells in gelatin methacrylate blended with laminin-411 and subsequent UV crosslinking (405 nm) resulted in a 3D-architecture encapsulating spatially distributed single cells (Appendix S2). After cultivation for 4 days, insulin-secreting cells in the bioprinted hydrogel started pseudoislet formation (Fig. 2A, Appendix S2). These multicellular aggregates grew over time in culture, and a diminishing number of remaining single cells permits the conclusion that cells are able to migrate within the hydrogel (Appendix S2). After long culture periods (>10 days) INS-1 pseudoislets could be observed to migrate out of the hydrogel structure due to the extensive pseudoislet density (not shown). The pseudoislets grew up to, but did not exceed, the size of average islets of Langerhans (150 μm; islet equivalency).^4, 14, 15^ For qualitative visualization of cell survival after printing and metabolic activity in 3D hydrogel culture, an MTT assay was used. Fig. 2A shows a sector of a 3D-bioprinted hydrogel grid with encapsulated, metabolically active pseudoislets. Quantitatively, the viability and proliferation of pseudoislet structures were measured with cytoplasmic ATP-based CellTiter-Glo^®^ 3D Viability Assay. Consistent with morphological observations, a significant 19-fold increase of luminescent signal was observed in the first 9 days and reached a plateau on day 12 (Fig. 2B). Proliferation of cells relies on sufficient oxygen and nutrient diffusion into the hydrogel structure to allow an entry into the cell cycle. Once maximum cellular density has been reached, pseudoislets start to migrate out of the hydrogel. Viability and proliferation of cells depends on, among other factors, the biomechanical properties and porosity of the encapsulating material. By application of different crosslinking times after printing, we showed that an optimum dose of UV light is required, as higher doses negatively impact the outcome (Fig. 2C-E). Even though 5 s crosslinking time per layer resulted in a hydrogel with viable pseudoislets, the number of islets is reduced compared with 2 s (Fig. 2C,E) A crosslinking time of 15 s resulted in a dense gel structure with no pseudoislet formation and a reduced number of INS-1 cells at the end of the experiment (Fig. 2D).

**Figure 2:**
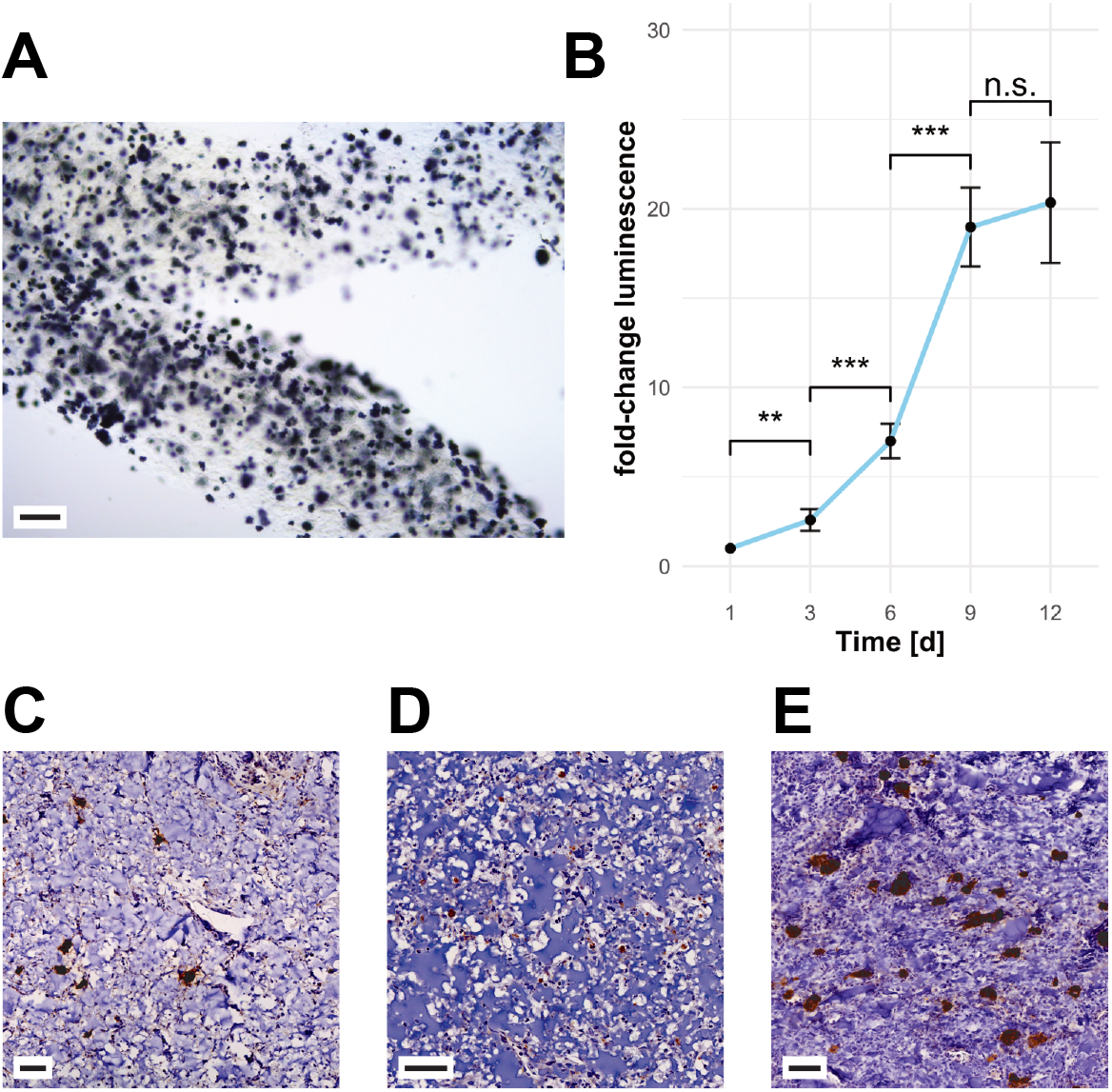
Hydrogel structure initiates pseudoislet formation. 3D-bioprinted INS-1 cells remain viable and proliferate. (A) MTT assay of pseudoislets formed after bioprinting (day 5 post-printing) of INS-1 containing gelatin methacrylate hydrogel shows metabolically active pseudoislets spatially distributed in 3D matrix. (B) Proliferation assay of bioprinted cells. A plateau is reached after 9 days in culture, indicating maximum loading capacity. Error bars depict SEM, n ≥ 13/timepoint. (C) CAM assay explant of bioprinted INS-containing hydrogel after 5 s of UV curing, (D) 15 s of UV curing, and (E) 2 s of UV curing. Crosslinking time and thus hydrogel stiffness is a crucial factor for the migration and reorganization capability. INS-containing hydrogel with 2 s crosslinking time enabled pseudoislet formation. Spatially distributed pseudoislets remained viable and functional. A longer crosslinking period resulted in higher hydrogel density and seemed to inhibit migration and growth of cells. Scale bar (A) 200 μm, (D-F) 50 μm.

### Transcriptome analysis of 3D-bioprinted insulin-secreting cells reveals enrichment of proliferative, anti-apoptotic pathways, structural integrity, and β-cell function

For a comprehensive understanding of the structural and functional implications of 3D-bioprinted hydrogel building blocks including insulin-secreting cells, global gene expression analysis (total mRNA) was performed using next-generation sequencing. 3D-Bioprinted grid structures with integrated insulin-secreting cells (Appendix S3A) were compared with their counterparts grown in monolayer (Fig. 3A,B). Differential gene expression analysis and subsequent gene enrichment analysis revealed significant alterations of predefined hallmark pathways (Fig. 3A, C-F; for MA-plot, see Appendix S3B).^16^ Sixteen hallmark pathways, including the sets for ‘TNF alpha signaling via NFkB’, ‘hypoxia’, ‘TGFβ signaling’, and ‘pancreas β-cell’, were significantly upregulated, whereas 11 pathways, including sets for ‘adipogenesis’, ‘reactive oxygen species pathway’, and ‘oxidative phosphorylation’, were downregulated (Fig. 3A; full report in Appendix S3C). The results are quantified using normalized enrichment score (NES)- and false detection rate (FDR)-adjusted p-values with a cut-off of 5% for hallmark gene sets. IPA revealed 100 significantly affected canonical pathways (cut-off threshold *P* < .05) (Fig. 3B; full report in Appendix S3D). A graphical network summary of the major biological themes for exploratory purposes is displayed in Appendix S3E. With regard to the effects of 3D-bioprinted hydrogel culture and the accompanying microenvironment that led to pseudoislet formation, evaluation of the enrichment of the pancreas β-cell-specific gene set (NES 1.63, *P* < .01) is of special interest (Fig. 3C). An upregulation of processes including regulation of insulin gene expression, glucose transport and sensing, modulation of ATP-sensitive potassium channels, secretory processes, and β-cell-specific canonical transcription factors and promotors was found (e.g., *Slc2a2, Sur1, Kcnj11),* whereas no significant difference in insulin gene expression itself could be detected (see excerpt table of differential gene expression in Appendix S3F). β-Cell-specific pathways were unbundled using IPA with Ingenuity Knowledge Base as reference and revealed an activation of insulin secretion signaling (z-score 3.657, *P* < .01) and IGF-1 signaling (z-score 1.667, *P* < .05) and an inactivation of type 2 DM signaling (z-score −1.265, *P* < .01). In contrast to *Mafa* and *Neurod1, Pdx1* is downregulated compared with monolayer control. However, causal network analysis identified all three transcription factors to be activated master regulators (activation z-score 3.182, bias-corrected *P* < .0001) based on 32 downstream genes in the dataset. The significant overexpression of IGF/TGFβ signaling cascade-related genes puts *Pdx1* expression in relation. IGF/TGFβ signaling is an important proliferative and functional regulator of β-cell growth and proliferation, and the enrichment data correlate with our findings in growth assays (TGFβ signaling NES 1.74, *P* < .01; Fig. 3E).^17^ Prolonged hyperglycemia induces glucotoxicity.^18^ An accumulation of glucose in 3D hydrogel culture may have led to GSK3B-regulated PDX1 phosphorylation and thus faster proteasomal degradation. High glucose induces FOXO1, which has been described to integrate β-cell proliferation with adaptive function to maintain tissue homeostasis at the center of IGF/TGFβ signaling.^17, 19^ Prolonged hyperglycemia may have triggered a transient inflammation of insulin-secreting cells, in which TGFβ1 interacts in the NFkB pathway (TNFA signaling via NFkB NES 2.37, *P* < .01).^17^ Additionally, overexpression of *Irs2* and *Atf3* has previously been reported to have a protective effect on β-cells under metabolic demand and alleviate hyperglycemia-induced apoptosis in support of our hypothesis.^20^ We hypothesize that INS-1 cells exposed to environmental stress due to the bioprinting process, transient hypoxic conditions, and hyperglycemia may have activated HIF1a/PFKFB3 stress-repair signaling.^21, 22^ This transient state is an adaptive, protective metabolic response that slows β-cell death at the expense of β-cell function and activates glycolysis (z-score 2.333, *P* < .0001) while inhibiting the oxidative phosphorylation (NES - 2.90, *P* < 0.01).^21, 22^ Additional upstream regulator analysis supported the likelihood of our hypothesis identifying glucose (activation z-score 3.835, overlap *P* < .0001) based on expression of 87 genes in the dataset (see the graphical display of glucose upstream regulator network incl. exemplary pathway overlay in Appendix S3G). Further, pseudoislet formation in 3D-bioprinted hydrogel culture led to a significant decline of pro-apoptotic genes *Bax*, *Bad*, and caspase 3 and overall activation of the functional annotation ‘Cell Viability’ (z-score 2.160, *P* < .0001) in IPA.

**Figure 3:**
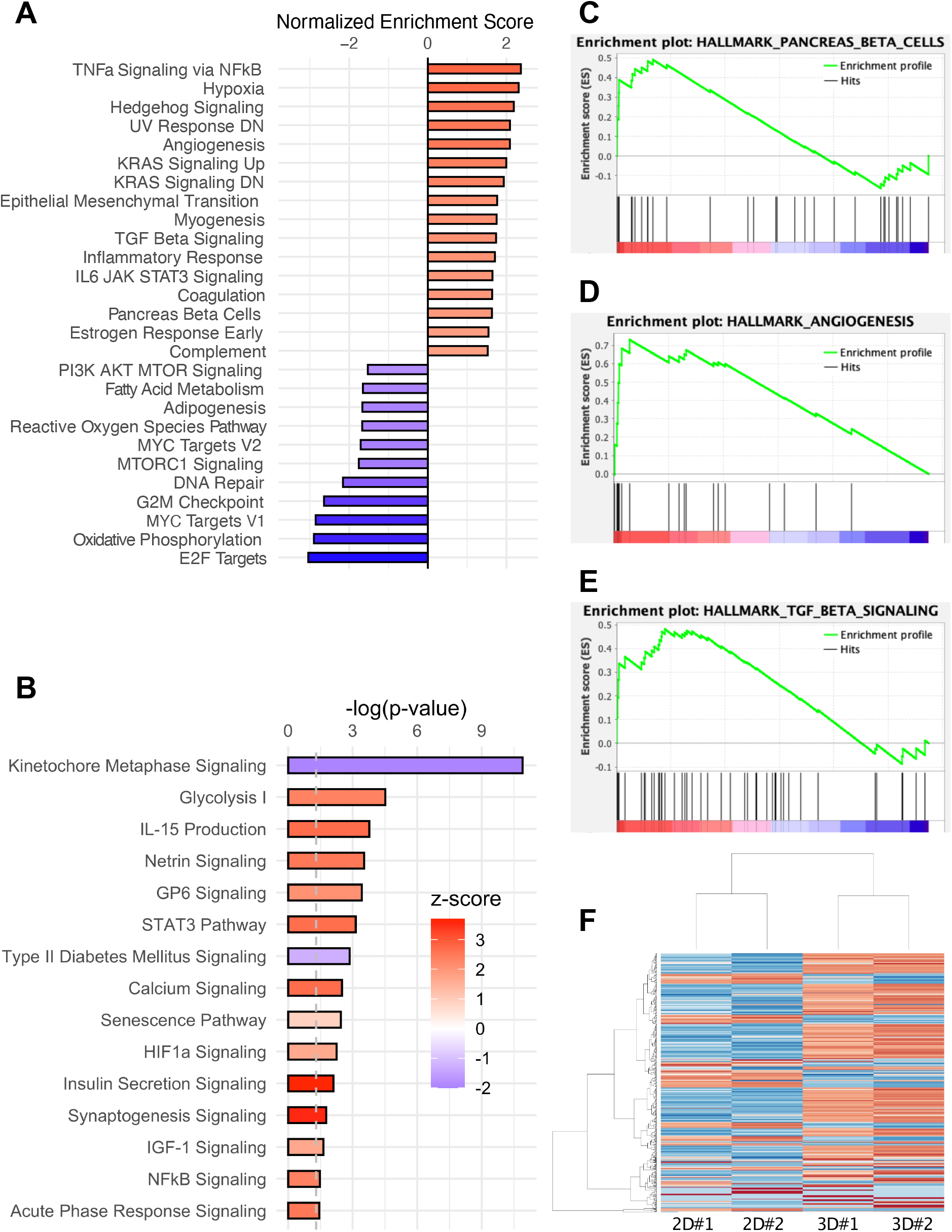
mRNA sequencing of 3D-bioprinted domes compared with monolayer culture control shows robust differential gene expression clusters and alteration of hallmark pathways. (A) Gene set enrichment analysis revealed significantly altered hallmark pathways (P < .05). (B) IPA revealed significantly altered canonical pathways (P < .05) and color-coded z-score indicates activation of pathways including insulin secretion signaling. Upregulated hallmark pathways include pancreas β-cells (C), TGFβ signaling (D), and angiogenesis (E). (F) Heatmap of alteration in gene enrichment (500 most significant alterations) comparing 3D-bioprinted culture with monolayer culture.

The above-described pseudoislet formation and proliferation in culture, together with reduced pro-apoptotic gene expression, may be a result of reduced anoikis due to cell-cell and cell-matrix interactions and anchorage-dependent growth. In islets, laminin-411 and laminin-511 are expressed and have been suggested to play an important role in β-cell proliferation and insulin transcription.^23, 24^ A hydrogel blend containing laminin-411 was therefore used in this study. The transcriptome data revealed significant overexpression of fibronectin, E-cadherin, basal cell adhesion molecules, and laminins secreted by INS-1 and known to be essential for structural integrity and cell contacts of islets, thereby enhancing functionality.^23-25^ In addition, significant overexpression of extracellular matrix (ECM)-localized growth factors such as *Vegfa* was found in INS-1 cells in 3D culture.

### Xenotransplantation to fertilized chicken eggs results in extensive vascular ingrowth and neoangiogenesis in scaffold components

Vascularization of building blocks is crucial for the viability and function of islets.^5, 9, 10, 26^ The transplantation of the PCL scaffold (Fig. 4A-C) and hydrogel (Fig. 4D-F) to the CAM of fertilized chicken eggs was utilized to investigate vascular ingrowth and vessel penetration through the solid PCL channel architecture in vivo (Fig. 4, Appendix S4A). The surface-functionalized PCL had beneficial effects on vascularization, as shown by the significantly enhanced total vessel network length (untreated control: 26856±3502SD [px], Hep-PCL: 34766±1650SD [px], *P* < .05) (Appendix S4A). Especially in the initial period after implantation, cells are faced with hypoxia and diffusion-based supply. We showed that insulin-secreting pseudoislets cultured in hydrogel structure survived this initial period before formation of vascularization. After 9 days of *in ovo* cultivation (Appendix S4B), the xenografts were explanted. A dense host-derived vascular network surrounding the xenograft was observed (Fig. 4E). Stereomicroscopy showed that some vessels penetrated the xenograft (Fig. 4F). Subsequent immunohistochemical analysis proved not only extensive peri-islet vascularization but also intra-islet vascularization (Fig. 4G,H). Staining with avian anti-CD34 showed capillary sprouting at the periphery and migration of host EC towards the graft center without additional external mitogenic stimulation (Appendix S4C). A direct comparison of heparin-conjugated and control scaffolds was possible in *ex ovo* CAM assay, where an anticoagulation effect of the heparinized scaffolds was observed (Appendix S4B, EDD11/18).

**Figure 4:**
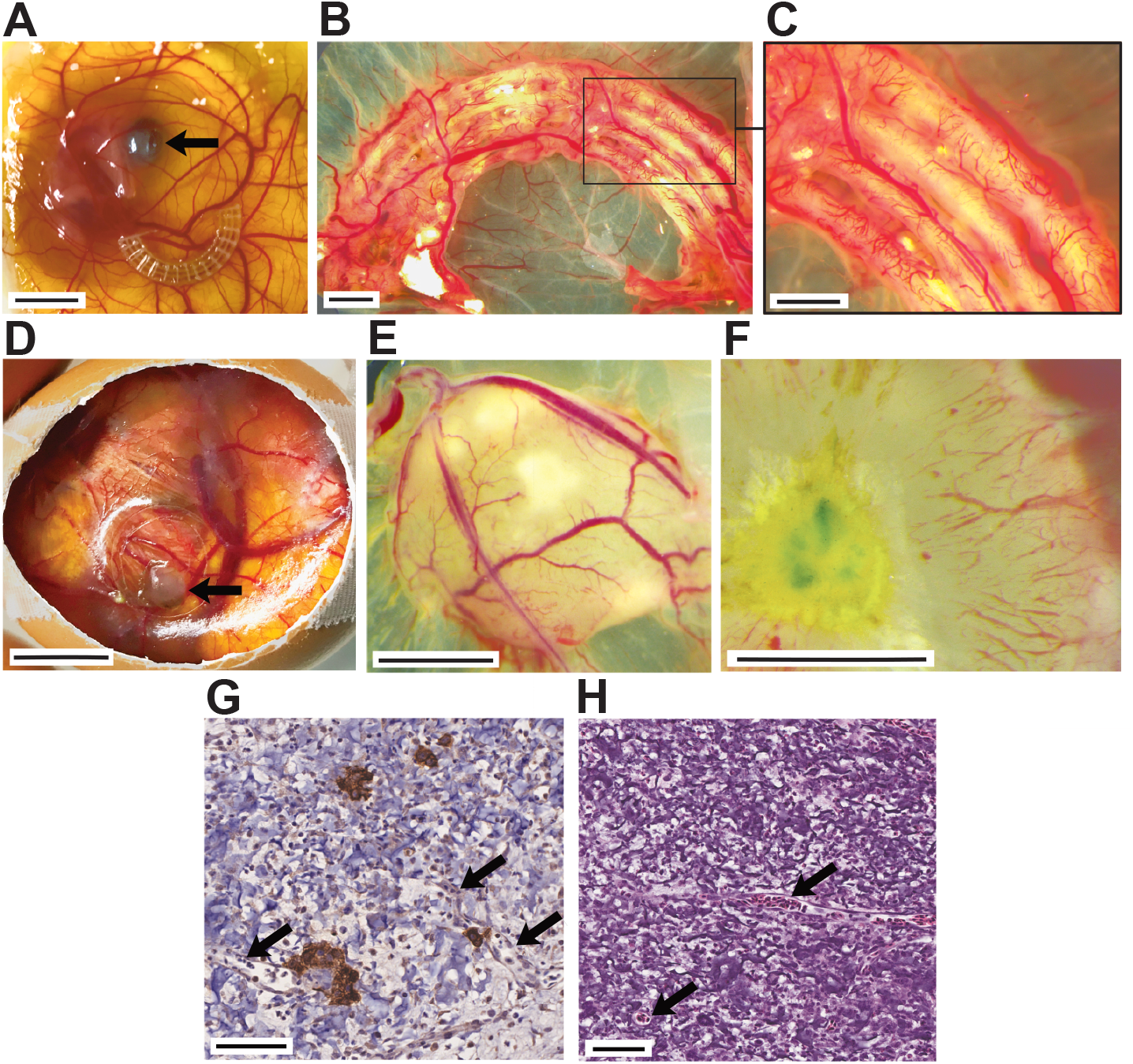
Chorioallantoic membrane assay is a suitable model for investigating angiogenesis in tissue-engineered grafts. Extensive, rapid vascular ingrowth is seen in both PCL and cell-laden hydrogel structure after the 9-day assay period. Ex ovo CAM assay experiments enabled direct comparison of heparinized PCL scaffolds with untreated controls and validated the beneficial properties of heparinization for enhanced vascular ingrowth (A-C, A: arrow indicates eye of chicken embryo). In ovo CAM assay experiments were used for investigation of 3D-bioprinted INS-1-laden droplets (D-F, D: arrow indicates xenotransplant). Vascular structures (arrows) penetrated into the scaffold (G, H). (G) Anti-insulin immunohistochemical staining of CAM assay explant. (H) H&E staining of CAM assay explant. Rapid vascularization maintained viability and function of pseudoislets. Peri- and intra-insular (asterisk) vessels were detected (G, H). Scale bar (A, D) 10 mm, (B, C, E, F) 2 mm, (G, H) 50 μm.

### Survival, viability, and insulin secretion of pseudoislets *in ovo*

3D-Bioprinted INS-1 cells formed pseudoislets *in vitro.* Pseudoislet formation was also observed after excision of *in ovo* cultured bioprinted hydrogel droplets. Xenografts remained on the CAM for 9 days supplied with nutrients only by diffusional processes from the CAM and newly formed vascular structures penetrating the xenograft. The CAM assay modeled the initial period after implantation, in which comprehensive vascularization of the graft is yet to develop. Insulin-secreting pseudoislets survived this initial period. Immunohistochemical staining against insulin demonstrated that pseudoislets remained functional until explantation (Fig. 2E, 4G). However, only pseudoislets with either minimal diffusion distance to the surrounding tissue (max. 650 μm; Appendix S5) or contact to intra-graft vessels (Fig. 4G) were found, therewith decreasing the homogeneity of the cellular distribution. Pseudoislets located deep inside the hydrogel structure experienced a hypoxic environment. Over a period of 9 days *in ovo*, anaerobic respiration was presumably insufficient to meet metabolic demand. Dependence on diffusion of nutrients and gases, soluble factors, and waste products from surrounding tissue into the graft limits the size if cell viability is to be retained. Homogeneous spatial distribution of pseudoislets in cross-sections of the graft was observed both at 20 μm and 40 μm distance from the bottom (Appendix S5). The graft area in these cross-sections of about 1.3 mm^2^ and 1.8 mm^2^, respectively, and the relatively small diffusion distance from graft base to CAM resulted in sufficient viability. At 60 μm diffusion distance, pseudoislets were predominantly located in marginal areas of the graft cross-section (6.45 mm^2^; Appendix S5). In addition, it is notable that pseudoislets are reduced in size with increasing diffusion distance. Pixel classification segmentation detected 1.8%, 2.1%, and 2.1% of insulin^+^ area at 20, 40, 60 μm cross-section layers, respectively. Examination of explanted specimens in which either the bioprinted graft had been UV-cured for a longer period (15 s) or a larger graft structure had been printed confirms the findings stated above (Appendix S5).

### Co-culture with EC ameliorates insulin secretion

The effect of the extracellular microenvironment and cellular crosstalk on the functional outcome was investigated by means of GSIS (Appendix S6). The 3D-bioprinted INS-1 group responded to glucose stimulation by secretion of 10.8 pmol/L and 21.4 pmol/L insulin in basal and high glucose concentration, respectively (*P* < .001, Fig. 5A). The functional outcome shows that cell encapsulation, bioprinting, and UV-crosslinking did not prevent the insulin-secretory function of bioink droplets (INS-1 group). The 3D-bioprinted co-culture group with HUVEC (INS-1/HUVEC, 1:2 ratio) yielded a secretion of 16.4 pmol/L and 31.7 pmol/L (*P* < .001, Fig. 5A). At both basal and high glucose concentrations, the coculture group showed significantly enhanced secretion of insulin (*P* < .05, *P* < .01). Both groups showed significantly increased insulin secretion after high-glucose treatment compared with basal glucose treatment (*P* < .001). Interestingly, the 2D-monolayer culture showed higher absolute insulin secretion after glucose challenge, possibly due to the tumor origin of the 2D adhesive cell line, a higher surface to volume ratio, and enhanced stimulus conduction of confluent monolayer cells (Appendix S6). However, the stimulation indices (high glucose/basal glucose ratio) were increased in both 3D culture conditions (1.56 [2D INS-1] compared with 1.97 [INS-1] and 1.93 [INS-1/HUVEC]). The stimulation in both 3D culture groups is similar to the characteristic index of healthy and freshly isolated human islets and can be attributed to the 3D matrix microenvironment.^27^

**Figure 5:**
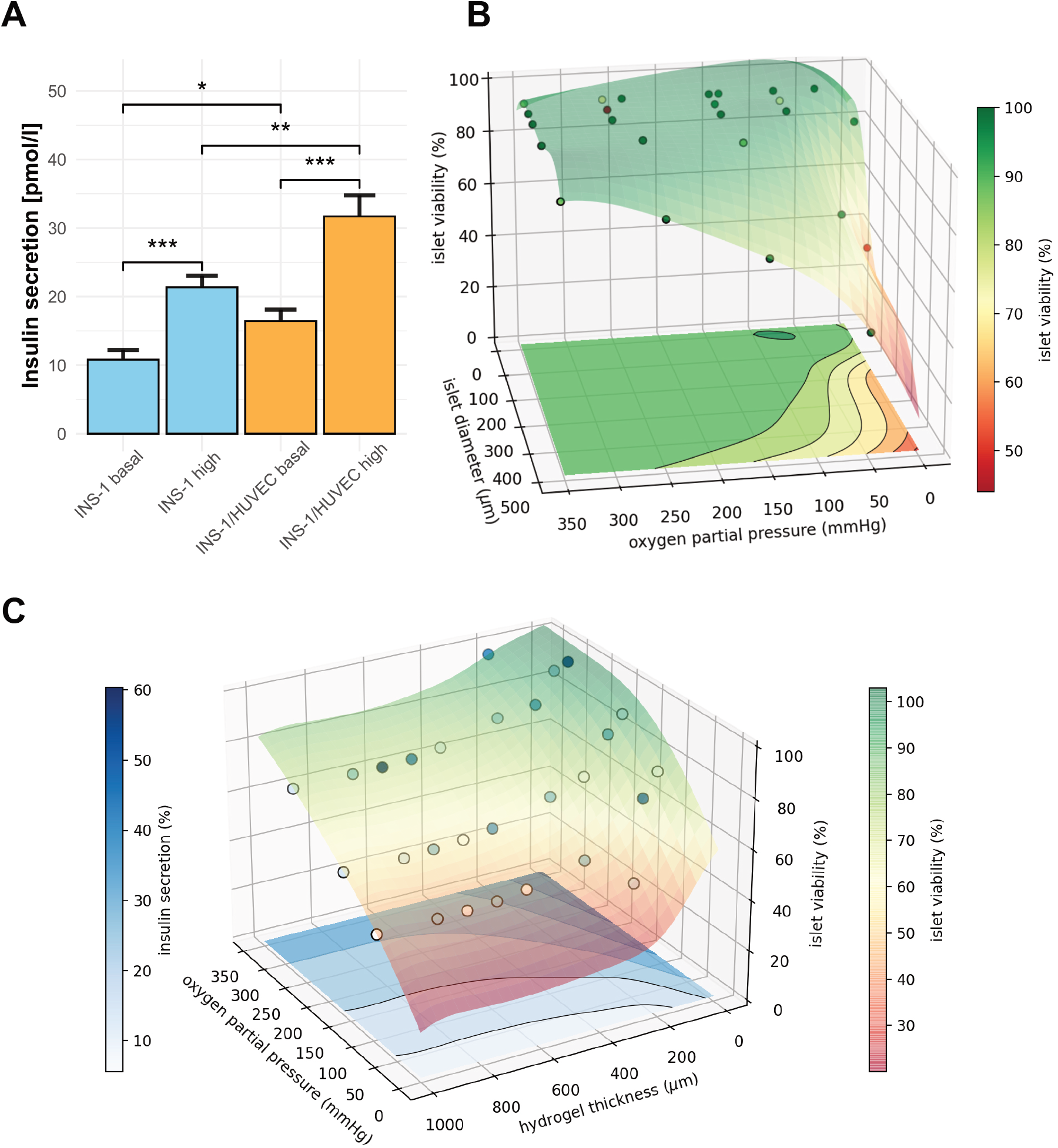
Insulin-secretion of bioprinted INS-1 cells is enhanced in co-culture with EC. Computer simulation of human Langerhans islets predicts concept feasibility and defines boundary conditions for viability and function in bioprinted 3D hydrogel. (A) Insulin secretion of hydrogel-embedded INS-1 culture and INS-1/HUVEC co-culture at basal and high glucose levels 3 days post printing. 3D-Bioprinted, encapsulated INS-1 cells are responsive to glucose stimulation. The INS-1/HUVEC coculture ameliorates the amount of insulin secreted. Insulin levels for both experimental settings were normalized to 10\000 cells. Error bars depict SEM, n ≥ 20/condition. (B) Simulation of viability of human islets of Langerhans encapsuled in hydrogel by finite element analysis. Calculation for 400 μm constant diffusion distance through the hydrogel. Glucose inflow concentration is kept at 10 mM. Inflow oxygen partial pressure ranging from 5 mmHg to 350 mmHg, islet diameter from 100 μm to 500 μm. Circles represent data points, gathered by diffusion simulation, and colored semitransparent surface and projection on the base represent fit through cell viability data. (C) Simulation of insulin secretion and viability. Constant glucose concentration of 10 mM, constant islet diameter of 500 μm, inflow oxygen partial pressure varying between 50 mmHg and 350 mmHg, hydrogel shell thickness ranging from 0 μm to 1000 μm. Circles represent calculated datapoints, semitransparent surface represents fit through 3D cell viability (right color bar). Insulin secretion is displayed as contour lines at the bottom of the diagram (left color bar). Simulations were performed for 464 different scenarios.

### Computer-aided applicability screening of scaffold architecture by finite element analysis demonstrates feasibility for Langerhans islets

The use of INS-1 cells instead of human islets could be viewed as a shortcoming of this study. Especially in terms of oxygen and nutrient diffusion the variable size of human islets^4^ needs to be considered to avoid function loss or central core necrosis.^3^ We addressed this requirement by performing simulations of diffusion gradients and metabolic activity of human islets. The intention was to define boundary conditions of scaffold architecture applicability, thus avoiding iterative experimental designs from bench to bedside. A primary limitation in translation of all tissue engineering approaches, in particular macrodevice scaffolds, is vascularization.^4^ According to previous findings, the diameter of the hydrogel capsule must extend no more than a few hundred micrometers in order to maintain the pseudoislet viability.^28^ The applicability of the scaffold architecture was validated by *in silico* modeling based on literature data^13^ (parameters displayed in Appendix S7). Finite element simulations with multiple input parameters predicted viability and insulin secretion of bioprinted islets of Langerhans as function of oxygen partial pressure, surrounding hydrogel thickness, and the diameter of the islet itself. Our simulations focused on the critical initial period after implantation, in which vascular ingrowth has not yet occurred. Hence, only diffusion processes were considered. Hypoxic conditions result in decreased viability and function, especially for large islet structures (Fig. 5B,C, Tab.1). Oxygen and glucose diffusion through islets is hampered compared with hydrogel diffusion (Appendix S7). Thus, large islets are more severely affected by hypoxia. As concluded before^13, 14^, small to average-sized islets containing bioink xenografts are more likely to survive. To a lesser yet considerable degree, cell viability with respect to oxygen levels is reduced with increasing hydrogel diffusion distance (Fig. 5C, Tab. 1). Finite element simulation supports identifying the best fit for bioprinting geometry. Insulin secretion by islets decreases rapidly with declining oxygen partial pressure and is diminished before viability begins to decrease. With a more than four times smaller diffusion coefficient, glucose diffusion is a more limiting factor for insulin secretion. However, simulations with hyperglycemic glucose concentrations showed decreased viability and function of islets (Tab. 1). High glucose concentrations stimulate insulin secretion in islet β-cells, leading to increased oxygen consumption. β-Cells in the islet shell consume more oxygen in response to hyperglycemia, leaving a smaller amount for core cells. Thus, islet core oxygen levels fall below a critical boundary for cell viability and cause core necrosis. The simulated functional outcome is additionally dependent on hydrogel thickness for insulin outflow. It is important to consider that due to the larger size of insulin, the overall secretion from the xenograft is also limited by accumulation and degradation of the hormone in the bioprinted hydrogel. The pseudoislets generated in this study and islet-like organoids generated from pluripotent stem cells^29^ range up to 150 μm in diameter and thus present more suitable prerequisites for diffusional nutrient supply while encapsulated.^15^

**Table 1:** Finite element analysis to prove feasibility for human Langerhans islets (excerpt). Anticipated results for cell viability and insulin secretion as function of oxygen level, surrounding glucose concentration, thickness of hydrogel encapsulation, and islet diameter.

| Oxygen level [mmHg] | Glucose concentration (initial) [mM] | Hydrogel thickness [$\mu\text{m}$] | Islet diameter [$\mu\text{m}$] | Viability [%] | Insulin secretion [%] |
| --- | --- | --- | --- | --- | --- |
| 160 | 10 | 50 | 500 | 92.1 | 46.53 |
| 160 | 10 | 50 | 150 | 100 | 70.24 |
| 90 | 10 | 50 | 500 | 78.5 | 32.46 |
| 90 | 10 | 50 | 150 | 100 | 70.24 |
| 90 | 5 | 50 | 500 | 87.2 | 14.10 |
| 90 | 25 | 50 | 500 | 73.8 | 41.26 |
| 90 | 5 | 500 | 500 | 54 | 3.82 |
| 90 | 25 | 500 | 500 | 40.3 | 8.25 |
| 90 | 5 | 1000 | 500 | 51.2 | 3.12 |
| 90 | 25 | 1000 | 500 | 37.5 | 6.2 |
| 90 | 5 | 50 | 150 | 100 | 28.76 |
| 90 | 25 | 50 | 150 | 100 | 95.96 |
| 90 | 5 | 500 | 150 | 100 | 27.61 |
| 90 | 25 | 500 | 150 | 100 | 69.56 |
| 90 | 5 | 1000 | 150 | 100 | 27.6 |
| 90 | 25 | 1000 | 150 | 99.8 | 64.85 |

## Discussion

In the present study, we developed and evaluated a concept for 3D-(bio)printing of insulin-secreting tissue as a novel treatment option for patients with insufficient insulin-secretory function. We proved the feasibility of our concept with regard to hybrid scaffold fabrication, integration of cells, and functional evaluation *in silico*, *in vitro*, and *in vivo*.

A hybrid scaffold was fabricated by 3D-(bio)printing using PCL and gelatin methacrylate hydrogel. Solid PCL as retrievable outer shell is a logistic template for a hybrid macrodevice. PCL is FDA-approved as a drug delivery device and is used in medical devices because of its biocompatibility, low immunogenicity, low foreign body response, and physicochemical properties.^11^ In this model, we propose using a permeable outer shell of a PCL mesh to support the inner insulin-secreting hydrogel core until it can support itself. Covalent heparin binding on the PCL surface improved vascularization *in vivo,* in agreement with a previous study by Marchioli et al.^7^, and increased cell adhesion due to decreased hydrophobic properties, increased protein binding capacity, and a different surface topography. EC coating of polymers has been reported to accelerate vascularization.^6^

Cellular integration was investigated by means of morphology, viability, proliferation, and transcriptome alterations. The bioprinted, multicellular hydrogel provides a microenvironment for insulin-secreting cells to form islet-like structures, thrive, and function. Total mRNA sequencing revealed that morphological pseudoislet formation in 3D culture correlates to upregulation of β-cell-specific proliferative pathways, insulin secretion signaling cascades, and angiogenesis pathways. The complex regulation and interaction of β-cell-enriched genes impedes a functional assessment of 3D-bioprinted hydrogel culture by transcriptome analysis alone. β-cell dedifferentiation after prolonged hyperglycemic conditions is a key mechanism causing a loss of functional β-cell mass in type 2 DM to alleviate apoptosis rates.^18^ However, we did not detect a functional consequence; dedifferentiation seems to be absent due to the overexpression of genes specific for the β-cell phenotype and robust activation of β-cell-specific pathways. This interpretation correlates with the findings from growth assays and stimulation index in GSIS experiments. Instead, the pseudoislets in 3D culture seem to respond to the metabolic demand by compensatory upregulation of *Foxo1, Iapp, Irs2,* and *Atf3* expression. VEGF-A, synthesized by β-cells among pancreatic cell types, is a crucial factor for communication with EC, and VEGF-A-depleted islets are only inefficiently vascularized when transplanted into a host compared with wildtype control.^23, 26^ In addition, islets depleted of *Vegfa* have a reduced number of capillaries and exhibit several defects in β-cell function, including insulin transcription, insulin content, first-phase insulin secretion, glucose tolerance, and β-cell proliferation.^23, 26^ In EC, VEGF-A will induce cell migration, proliferation, maintenance of capillary fenestrations, and β-cell mass. ^3, 5, 23^ During clinical islet isolation the dense capillary network becomes disrupted.^5, 15^ The capillary network or EC networks play an essential role in vascularization and thus survival of islets after transplantation.^5-7, 15^ However, previous studies have shown a concentration-dependent adverse effect of additional VEGF-A.^7^ Our results show significantly upregulated expression of VEGF-A after cell integration in the hydrogel structure and extensive vascular penetration and neoangiogenesis *in ovo*. The microenvironmental gradient of VEGF-A may already have reached a therapeutic gradient without external VEGF-A delivery. Our findings support omitting obligatory additional VEGF-A delivery in building blocks, as overdosage will result in leaky and dysfunctional vessels.^5, 9^

Functional evaluation of insulin secretion after glucose challenge showed the superiority of co-culture with EC in providing a natural cellular niche. The beneficial effect of EC on secretory function of insulin-secreting cells has been reported before.^5, 23, 24^ As an essential element of pancreatic vessels, EC contribute to the delivery of glucose as the primary input of β-cell signaling and transport insulin as functional output to its target location. In addition, direct and paracrine communication between β-cells and EC enhances the structural and functional integrity of islets and their insulin-secretory function.^5, 24^ Thrombospondin-1, endothelin-1, and hepatocyte growth factor secreted by EC may have directly stimulated insulin secretion.^5^ In addition, ECM secretion by HUVEC influences the β-cell microenvironment.^5, 24^ As reported previously, external delivery of integral basement membrane proteins can compensate only partially for ECM and EC loss during islet isolation^23^, and this explains the superior functional outcome of co-culture hydrogel compared with laminin-containing hydrogel with INS-1 alone. Due to the absence of an immune system in the developing chick embryo,^30^ CAM xenografts are not suited to assess interactions of the scaffold with the host immune system, including protection of insulin-secreting pseudoislets by hydrogel encapsulation itself. In the developmental stage used in the CAM assay, the chicken immune system is still developing and is not yet fully functional. Further investigations including *in vivo* trials are necessary. The feasibility of the concept for integration of human islets was evaluated by computer simulations using finite element analysis. Especially small and average-sized islets, similar to the pseudoislets formed by INS-1 cells, are feasible for integration into a bioprinting process, as hypoxic conditions in the core of larger islets reduce overall viability and function. Scaffold devices have often been implanted in the subcutaneous space, resulting in low potential for vascularization and lack of glycemic control.^8, 9^

In the overall concept, findings from this study are merged with findings from previously published evidence gap maps, showing that transplantation of scaffold-based constructs alongside or in close proximity to vascular structures has been rarely investigated.^3, 8^ We propose that attachment in close proximity to neurovascular structures will contribute to vessel formation while avoiding the instant blood-mediated inflammatory reactions that lead to graft loss, as described in clinical islet transplantation.^4^ The parametric design allows generation of patient-specific devices that can be customized depending on, for example, the specific anatomy, and overcomes scalability concerns. Theoretically, building blocks may be fused to form larger, multifunctional geometrical units augmenting the flexibility of tissue engineering applications. In the future, patients suffering from either type 1 or type 3c DM might benefit from this bioartificial insulin-secreting device by virtue of improved glycemic control, insulin independence, and quality of life. Although both these types of DM evolve through a loss of insulin-secreting cells, the cell sources used for transplantation can differ.^2^ Transplantation of autologous islets diminishes immunological reactions towards the graft.^2^ In the case of allogenic islets and other cell sources, however, more research on the immunoprotective capabilities of the device is required. If necessary, local immune modulation by functionalization of immunosuppressive agents as described earlier might be harnessed.^3^ Differentiation and integration of progenitor and stem cells as an unlimited cell source has the potential to expand the applicability of our concept and future bioconvergence research^2, 3^

## Conclusion

We have shown that 3D-(bio)printing is a feasible method for fabrication of a hybrid scaffold for insulin secretion. Integration of cells into the hydrogel structure initiates pseudoislet formation at the microscopic level and cell proliferation at the molecular level. Functional evaluation by means of insulin secretion of the bioprinted xenograft have been demonstrated *in vitro* and *in ovo*. Rapid vascular ingrowth and neoangiogenesis can be found in all scaffold components as a premise for long-term function *in vivo.* We have shown on the one hand that the functional properties of the scaffold itself can be tuned at multiple levels and on the other hand that multicellular composition can further contribute to mimicking the natural microenvironment, thus improving functional outcomes. We propose a modular, patient-specific device architecture for encasement of neurovascular structures. Future experiments with human islets will be necessary to verify our concept for future *in vivo* trials.

## Data availability statement

RNA sequencing datasets are publicly available in the GEO repository. The GEO accession number is GSE166285. Other datasets generated for this study are available from the corresponding authors on request.

## Acknowledgments

We express our gratitude to S. Le Blanc, Ph.D., K. Felix, Ph.D., L. Liu, Ph.D., D. Schmitt, M.Sc., and W. Wagner (all University Hospital Heidelberg, Germany) for their helpful discussion and to K. Schneider and K. Ruf of the European Pancreas Center for their excellent technical support. Electron microscopy was performed at the EM Core Facility, Heidelberg University, and the technical assistance of S. Wurzbacher is acknowledged. RNA sequencing was performed at the Genomics Core Facility of the European Molecular Biology Laboratory (EMBL) Heidelberg under the supervision of V. Benes, Ph.D. Translational Lung Research Center (TLRC) Heidelberg provided access to whole-slide scanning and fluorescence microscopy.

## Author contributions

Conceptualization: G.A.S., H.G.K.; methodology: G.A.S., I.H., N.A.G.; investigation: G.A.S., E.P., D.C.; validation: G.A.S., E.P.; formal analysis: G.A.S., E.P., M.N.M., D.C., C.B-P; software: V.V., D.C., C.B-P; data curation: G.A.S.; visualization: G.A.S., E.P., M.N.M., D.C.; writing - original draft: G.A.S.; writing - review and editing: E.P., M.N.M., D.C., F.N., I.H., N.A.G., T.H., H.G.K.; resources: F.N., I.H., N.A.G., T.H., H.G.K.; funding acquisition: G.A.S., H.G.K.; project administration: H.G.K.; supervision: T.H., H.G.K.

## Transcript profiling

The GEO accession number is GSE166285

