## SupplementaryMaterial for "Towards 3D-Bioprinting of an Endocrine Pancreas: A Building-Block Concept for Bioartificial Insulin-Secreting Tissue"

### Supplementary Materials

#### Materials and Methods – full report

**Computer-aided design (CAD) model creation and slicing for hybrid scaffold fabrication.** CAD models of the scaffold structures were created using the open source package Blender[1]. The models were converted from Standard Triangulation Language (STL) to numerical control G programming language using the Cura software package, v4.1 (Ultimaker, Utrecht, NL; available from https://www.ultimaker.com/en/products/ultimaker-cura-software) for dual-extrusion 3D printing. For 3D bioprinting an integrated slicing software was applied (CellInk, Gothenburg, Sweden).

**3D-Printing of PCL, heparin surface functionalization, growth factor addition for hybrid scaffold.** PCL components were fabricated using a dual-extrusion-based 3D printer (UM S5, Ultimaker, Utrecht, Netherlands). Polycaprolactone (PCL) filament (Facilan$^{TM}$ PCL 100 Filament 2.85 mm, 3D4Makers, Haarlem, Netherlands; MW: 50 000 g/mol) was used for the scaffold structure and polyvinyl alcohol filament (PVA; Ultimaker) as a sacrificial, water-soluble support structure. PCL was extruded with an AA 0.25 mm, PVA with a BB 0.4 mm print head, using the following settings: print speed 20 mm/s, build plate temperature 30 °C, fan speed 100%, AA print head temperature 140 °C, BB print head temperature 215 °C. In order to print PCL at temperatures as low as 140 °C, the g-code was manually edited by prefix code 'M302' to avoid device-specific conformity checks. For heparin surface functionalization, 1% (w/v) heparin (Lot# H0200000, Merck, Darmstadt, Germany) was dissolved in 0.05 m 2-(N-morpholino) ethanesulfonic acid monohydrate (MES) buffer (Lot# K49565026903, Merck) at a pH of 5.5. Quantities of 0.5 m 1-ethyl-3-(3-dimethylaminopropyl) carbodiimide (EDC) (Lot# E7750, Merck) and 0.5 m N-hydroxysuccinimide (NHS) (Lot# BCBW6640, Merck) were added to the heparin solution. Scaffolds were previously equilibrated for 30 min in MES buffer and subsequently immersed in reaction mixture. The reaction mixture was then stirred for 8 h at room temperature. The reaction was stopped by extensive washing with sterile $H_2O$ to remove unbound heparin. Growth factor addition was performed by immersion of scaffolds in beta fibroblast growth factor (bFGF; 500 ng/ml) or nerve growth factor (NGF; 500 ng/ml) in phosphate-buffered saline (PBS) for 2 h at room temperature. Scaffolds were stored in PBS. Dried PCL scaffolds, heparin-coated PCL scaffolds, and heparin-coated PCL scaffolds after growth factor addition were 10 nm gold/platinum sputtered (Leica EM ACE 600, Leica Microsystems GmbH, Wetzlar, Germany) and qualitatively analyzed by scanning electron microscopy (Zeiss Leo Gemini 1530, Carl Zeiss AG, Oberkochen, Germany). Images were taken at different magnifications with an accelerating voltage of 2.0 kV.

**Cell culture.** The rat INS-1 832/3 cell line (referred to as INS-1 hereinafter) was obtained from Merck (Darmstadt, Germany). A HUVEC cell line was obtained from the American Type Culture Collection (Manassas, VA, USA). Mycoplasma testing was performed monthly by polymerase cell reaction. INS-1 cells were used until passage 10; insulin-producing function was ensured by selection through Geneticin resistance. INS-1 cells were cultivated in RPMI-1640 (Gibco, Thermo Fisher Scientific, Waltham, MA, USA) supplemented with 10% fetal bovine serum (Gibco), 1% Geneticin (Merck), 1% HEPES 1M (Gibco), 1% sodium pyruvate 100 mM (Merck), and 0.1% 2-mercaptoethanol (Merck). HUVEC cells were cultivated in endothelial cell growth medium (Lot# 211-500, Cell Applications, San Diego, CA,

USA) supplemented with 1% penicillin/streptomycin (Merck) and 5% fetal bovine serum. For co-cultures, culture medium composition was chosen according to cell ratio. Cells were grown in T75 flasks (Falcon®, Corning, NY, USA) at 37 °C and 5% $CO_2$.

**Bioprinting of cell-laden hydrogels for hybrid scaffold.** Bioprinting was performed using the BioX from CellInk. Pneumatic extrusion print heads were used for extrusion of bioink. $3x10^6$ cells/ml hydrogel were used for bioprinting. The cells, either INS-1 only or INS-1 with HUVEC cells in 1:2 ratio, were diluted in either RPMI-1640 or a 1:2 mixture of RPMI-1640 and endothelial cell growth medium and gently mixed 1:10 with GelXA LAMININK-411 hydrogel (Lot# IK-3X2123, CellInk) using female-female Luer-lock-adapted syringes. The INS-1/HUVEC ratio was chosen based on the natural islet microenvironment[2] and, due to superior results, compared with a 1:5 ratio. The cell-laden hydrogel was transferred to a UV-shielded cartridge and centrifuged at 100 g for 1 min to remove any air. The cartridge (pre-cooled to 4 °C) was loaded into pneumatic print heads. Bioprinting in 24-well plates (Falcon®, Corning) was performed with the following settings for proliferation assays, CAM xenotransplantation, and glucose-stimulated insulin secretion (GSIS): droplet print mode, 2.6 s extrusion time, 30 kPa extrusion pressure, 2 s ultraviolet (UV) crosslinking (405 nm) at 5 cm distance. After printing, the hydrogel domes were incubated in 1 ml of either RPMI-1640 (INS-1 only) or in a 1:2 mixture RPMI-1640 and endothelial cell growth medium (co-culture). Grid-like structures were printed in 24-well plates to perform metabolic assays and total RNA isolation using a 21-gauge conical nozzle, extrusion pressure 23 kPa, print speed 8 mm/s, 50 ms pre-flow delay, infill 15%, 2 s crosslinking at 405 nm with 5 cm distance to printed layer.

**Detection of metabolic activity and proliferation.** For a visual assessment of metabolic activity, INS-1 cells in bioprinted grid scaffolds were stained with thiazolyl blue tetrazolium bromide (MTT, Merck) after 5 days in culture. A quantity of 100 µL of 5 mg/ml MTT dissolved in PBS was added to 900 µL INS-1 culture medium and scaffolds were incubated under cell culture conditions for 2 h. Images were taken using a Leica DMi8 fluorescent microscope. The viability of the bioprinted, encapsulated INS-1 cells was determined using an ATP-based assay with luminescent readout (CTG, CellTiter-Glo® 3D Cell Viability Assay, Promega GmbH, Walldorf, Germany) according to the manufacturer's protocol. In brief, droplets printed in 96-well plates were incubated with 100 µl INS-1 expansion medium. On days 0, 3, 6, 9, and 12 after printing, blinded sample droplets together with 100 µl medium were transferred into a 96-well solid white polystyrene microplate (Falcon®, Corning) and 100 µl CTG reagent was added to each well. The microplate was continuously shaken for 25 min, and luminescence was measured using an ELISA reader (Synergy HTX, multi-mode reader, BioTek, Bad Friedrichshall, Germany).

**RNA sequencing.** Genome-wide expression profiling was a service provided by the European Molecular Biology Laboratory (EMBL; Heidelberg, Germany). After 5 days in culture total RNA was isolated from 2D monolayer culture and 3D hydrogel culture using an RNeasy Mini kit (Qiagen, Hilden, Germany) according to the manufacturer's instructions (biological replicates, passage 3). After isolation, the total RNA was treated with the Turbo DNAfree kit according to the manufacturer's instructions (Thermo Fisher Scientific). The RNA concentration and quality were evaluated using Nanodrop and Agilent2000 Bioanalyzer (Appendix X). RNAseq libraries were prepared using the TruSeq stranded mRNA kit and sequenced using an Illumina NextSeq 500 platform, resulting in 75-bp single end reads in a read count of 36 million reads per sample. Quality control of the RNAseq FastQ files was performed with FastQC v.0.11.8. The obtained reads were pseudoaligned using the rn4 reference genome with the addition of human insulin gene and quantified by Salmon v1.2 with standard parameters. The resulting transcript expression levels were summarized to gene-level expression values and corrected for average transcript length by using tximport v1.10.1 and the "lengthScaledTPM" option while filtering out genes expressed in low amounts (average counts < 10).[3] Differentially expressed genes for the culture conditions were determined by using the DESeq2 v1.22.2 package.[4] Using the DESeq2 and log2 fold change pre-ranked differentially expressed genes, a gene set enrichment analysis was performed using the fgsea package v1.8 and the hallmark gene sets from MSigDB v7.1.[5] Additional data analysis was performed using Ingenuity Pathway Analysis (IPA; Ingenuity Systems, Qiagen) by input of gene identifiers, log2 fold change, and p-values.[6] Canonical pathway analysis identified the pathways referenced in the Ingenuity Knowledge Base of canonical pathways (11/2020) that were significant to the data set ($p \leq 0.05$). Molecules from the data set that met the log fold change cut-off of <-0.5 and >0.5 and a p-value ≤ 0.05 were considered for the analysis.

**Xenotransplantation to the chorioallantoic membrane (CAM) of fertilized chicken eggs.** As described before,[7] fertilized eggs from genetically identical hybrid Lohman Brown chickens were obtained from a local ecological hatchery (Gefluegelzucht Hockenberger, Eppingen, Germany). Eggs were delivered at day 0 of chick development and were immediately cleaned with 70% warm ethanol. The eggs were placed in a digital motor breeder (Type 168/D, Siepmann GmbH, Herdecke, Germany) at 37.8 °C and 45–55% humidity with an activated turning mechanism to start day 1 of the embryonic chick development. Four days after incubation, the turning mechanism of the incubator was switched off and a small hole was cut into the eggshell to detach the embryonic structures from the eggshell by removing 3 ml albumin. The hole was covered with Leukosilk® tape (BSN medical, Hamburg, Germany), and the eggs were incubated further with the turning mechanism switched off. On day 9 of embryonic development, the tape was removed and the epithelial layer of the chorioallantoic membrane (CAM) was gently scratched with a syringe needle to ensure immediate blood supply to the xenotransplant/polymer component. PCL scaffold groups and bioprinted xenografts (bioprinted hydrogel) were placed on the CAM. PCL scaffold groups consisted of 3D-printed PCL scaffolds functionalized with covalently bound heparin and plain PCL scaffolds. Prior to implantation, scaffolds were sterilized with 70% ethanol for 48 h. For explantation, the chicks were ethically euthanized at day 18 of development, 3 days before hatching, as described before.[8] PCL scaffolds and bioprinted xenografts were excised including the surrounding CAM and briefly washed in PBS before further imaging. Each specimen was imaged by stereomicroscopy (Leica MZ10 F, Leica Microsystems GmbH, Wetzlar, Germany). Images of PCL scaffold groups were analyzed using an automatic image analysis software (WimCAM; CAM Assay Image Analysis Solution, Release 1.1, Wimasis, 2016).

**Immunohistochemistry of xenograft tissue.** Xenografts were fixated in 5% formaldehyde (Otto Fischar GmbH & Co. KG, Saarbruecken, Germany) after excision and transferred to 70% ethanol after 24 h. The fixated, explanted xenografts were embedded using HistoGel™ (Lot# 370234, HG-4000-012, Thermo Fisher Scientific) and cryomolds (Tissue-Tek™, Cryomold™, Thermo Fisher Scientific) according to the manufacturers' instructions. After paraffin embedding of the xenografts, randomly chosen blocks from each group were continuously sampled in 5 µm serial sections, numbered, and processed for histology. Slides with odd numbers were stained with Mayer's Hematoxylin-Eosin (H/E), while those with even numbers were immunostained for insulin. Therefore, a primary anti-insulin antibody (monoclonal mouse IgG, 2D11-H5, Lot# SC-8033, SantaCruz, Dallas, TX, USA), overnight 1:100 in background reducing antibody diluent (S3022, Dako, Agilent Tech., Santa Clara, CA, USA), and a polyclonal goat anti-mouse secondary antibody (Dako, Agilent Tech.), 3-3'diaminobenzidine staining with subsequent hematoxylin counter-staining, were used. In addition, randomly chosen samples were immunostained for endothelial and endothelial progenitor cells with a primary anti-chicken CD34 antibody (monoclonal mouse IgG; Lot# AV138, UniProt E1BUT3, Avian Immunology toolbox project, Bio-Rad Laboratories GmbH, Feldkirchen, Germany) to identify newly formed vascular structures in the CAM assay. Whole slides were scanned at 40× magnification using a NanoZoomer S60 Digital Slide Scanner (Hamamatsu Photonics, Hamamatsu City, Japan). Stained tissue slides were analyzed using the ilastik software package[9] for supervised machine learning (ilastik: interactive machine learning for [bio]image analysis, v1.3.3, open-source, https://www.ilastik.org/download.html). Islets were segmented using the pixel classification workflow (islet, non-islet, background [not islet, not non-islet]). First, a random forest classifier was trained manually, and subsequent batch processing was performed. Due to limitations of the machine-learning strategy in differentiating xenograft and CAM tissue, the xenograft area was determined using ImageJ (Fiji package[10]).

**Glucose-stimulated insulin secretion (GSIS).** For GSIS experiments, INS-1 cells were stained with red fluorescent membrane inserting dye PKH-26 (Lot# SLBW0232, Merck) according to the manufacturer's protocol prior to mixing with the hydrogel for bioprinting. In brief, cells were trypsinized using 0.25% Trypsin-EDTA (Gibco), rinsed with Dulbecco's PBS (DPBS; PromoCell GmbH, Heidelberg, Germany), and finally pelleted. The pellet was resuspended in Diluent A, and PKH-26 dye dissolved in Diluent A was added to the cells. After rapid mixing and incubation, culture medium was added. The cell suspension was centrifuged and further washing steps were performed. The insulin secretion of 3D-bioprinted INS-1 (low glucose: n = 22; high glucose: n = 20) and INS-1/HUVEC co-culture (low glucose: n = 22; high glucose: n = 21) groups, INS-1 cells seeded on PCL/heparin-PCL scaffolds ($2x10^5$ cells in 1ml RPMI-1640 in per well), and the 2D monolayer control group was measured. In the 2D monolayer culture group, INS-1 cells were seeded in 4-well chamber slides ($10^5$ cells in 1 ml RPMI-1640 per well) (Nunc® Lab-Tek®, Thermo Fisher Scientific). The medium was changed after 2 days, and GSIS was performed on day 3 in all conditions. For preparation of the GSIS solution, SILAC RPMI-1640 Flex (A2494201, Gibco) was supplemented with $MgSO_4$ (1.16 mmol/l end concentration) (Merck), $CaCl_2$ (2.5 mmol/l end concentration) (Merck), 20 mM HEPES, and 0.2% BSA (Merck). GSIS was initiated by rinsing the cells once with low-glucose solution (1.67 mM D-glucose), followed by incubation for 1 h in 1 ml low-glucose solution. After that, either 1 ml of low-glucose solution or 1 ml of high glucose solution (16.7 mM D-glucose) was added, followed by incubation for 2 h. A quantity of 500 µl medium was taken and briefly spun down in a 1.5-ml Eppendorf tube. Next, 400 µl supernatant was used for determination of insulin concentration by chemiluminescence immunoassay (ADVIA CENTAUR, Siemens Medical Solutions, Malvern, PA, USA). After GSIS of 2D samples on chamber slides, cells were incubated in 5% formaldehyde solution for 15 min, rinsed with DPBS twice, dried for 10 min, and covered with Fluoroshield Mounting Medium with 4′,6-diamidino-2-phenylindole (DAPI; Abcam, Cambridge, UK) and a coverslip. Similarly, 3D-bioprinted samples were fixated and transferred to a glass slide, covered with two drops of Shandon Consul mounting medium (Thermo Fisher Scientific), and squashed with a coverslip until flattened. Cells were counted using a Leica DMi8 fluorescence microscope with the following settings for PKH-26 imaging: 10× magnification, Y3 filter block, 260 ms exposure time, gain 7. DAPI imaging was performed with the following settings: 10× magnification, DAPI filter block, 10.5 ms exposure time, gain 4. Image processing was performed using Leica LAS X software, and subimages were assembled to mosaics depicting whole domes or whole well bottoms. Cells were counted using ImageJ (Fiji package[10]). In the case of polymer scaffold culture, cells were lysed using radioimmunoprecipitation assay (RIPA) buffer supplemented with protease inhibitor (cOmplete Mini, Roche, Basel, Switzerland) and incubated on ice for 10 min. Protein concentration was determined using a bicinchoninic acid (BCA) assay (Pierce BCA Protein Assay Kit, Thermo Fisher Scientific). The assay was performed according to the manufacturer's protocol. In n=12 wells of a 24-well plate, $10^5$ INS-1 cells were seeded in 1 ml RPMI-1640 for correlation of total protein to cell number. After 48 h the medium was changed, followed by another 24 h of incubation. Cells were lysed using 250 µl RIPA buffer + protease inhibitor in n=6 wells and total protein was determined. The residual wells were fixed with 5% formalin, rinsed twice with PBS, and mounted using Fluoroshield Mounting Medium with DAPI. After cell counting, a conversion factor between cell number and total protein was obtained.

**Computer-aided applicability screening of scaffold architecture by finite element analysis.** Diffusion of oxygen, glucose, and secreted insulin through islets of Langerhans encapsuled in a hydrogel shell was modeled using a custom python script (v3.8, Python Software Foundation, https://www.python.org) for input parameter-based insulin secretion based on literature data[11] (s. Appendix S7). The finite element simulations are based on mesh generation using the open-source module Gmsh[12] and FiPy[13], a respective finite element solver. The simulations were performed in 2D and the results were extrapolated to 3D spherical setups. Hydrogel shell and islet were initialized with 10 mM (5, 15, 25 mM) glucose. The initial oxygen partial pressures ranged from 90 mmHg to 270 mmHg. The thickness of the hydrogel shell varied between 0 µm and 1000 µM. In the simulation, diffusion started from outside the capsule and triggered consumption of glucose and oxygen within the islets. The simulations were carried out for at least 60 s with step sizes for diffusion below 0.005 s leading to converged results. Based on simulation results, cell viability was evaluated by considering a minimum local oxygen partial pressure of 0.07 mmHg for cells to survive.

**Statistical analysis.** Data analysis and statistical testing was done using R version 3.6.1 and ggplot2 package. Proliferation assay, vascular ingrowth analysis, and GSIS were analyzed by non-parametric Wilcoxon rank-sum test. Conditions in 2D and 3D GSIS were normalized to 10 000 cells, and PCL scaffold conditions were normalized to 100 µg total protein. Values not within the 2 ⌠ interval were classified as outliers and removed prior to analysis. The results are expressed as standard error of the mean. Sequencing was performed using two biological replicates; other experiments were repeated at least three times. Using IPA), the significance of association between dataset and canonical pathway was measured in two ways: (1) ratio of dataset molecule number mapped to pathway divided by total molecule number mapped to canonical pathway; (2) right-tailed Fisher's exact test to calculate a p-value determining the probability that the association between dataset genes and canonical pathway is explained by chance alone. Molecules from the data set that met the log fold change cut-off of <-0.5 and >0.5 and a p-value ≤ 0.05 were considered for the analysis. R version 4.0.0 with additional packages tidyverse v1.3, ggpubr v0.4, and ggrepel v0.8.2 (https://www.r-project.org) was used for data analysis and presentation. Statistical significance is depicted by means of asterisks. p-values are given as *$P \leq .05$; **$P \leq .01$, ***$P \leq .001$; ****$P \leq .0001$.

#### Appendix S1A

3D-printed polycaprolactone component

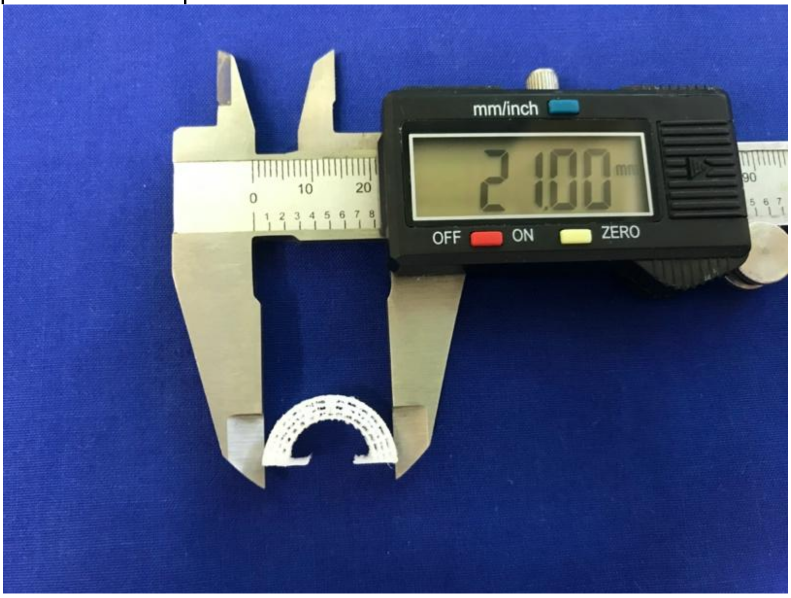


#### Appendix S1B

Glucose-stimulated insulin-secretion: INS-1 cells seeded on 3D-printed polymer scaffolds

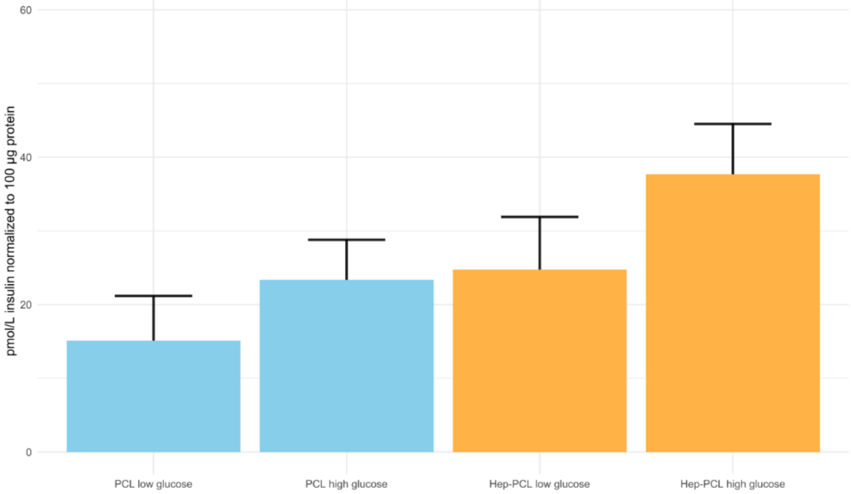

### Appendix S2

3D-bioprinted droplets: gelatin methacrylate blend / INS-1 cells

| | |
|---|---|
| 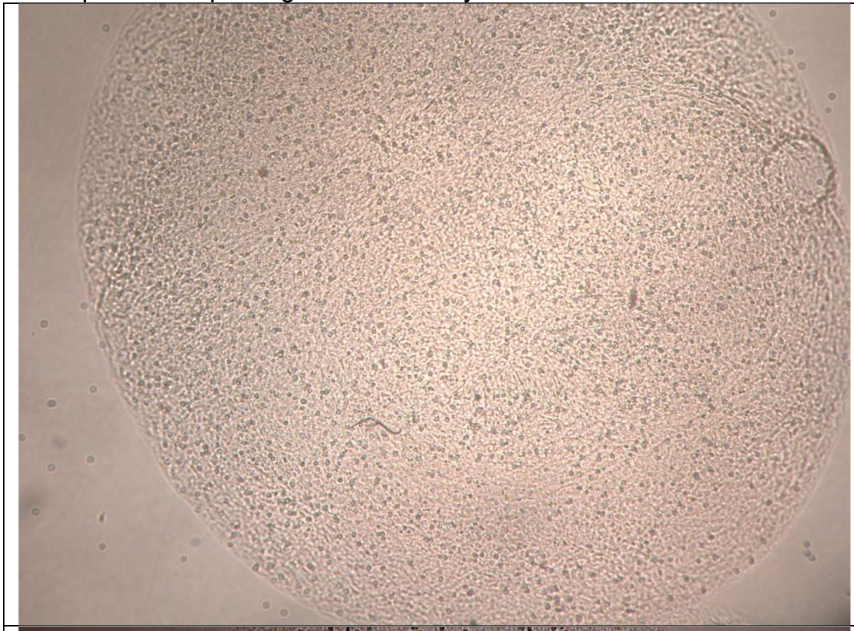 | Day 1 post printing<br><br>INS-1 832/3 cells<br>CellInk GelXA LAMININK 411<br>Seeding density<br>2×10^6/ml<br><br>4× magnification |
| 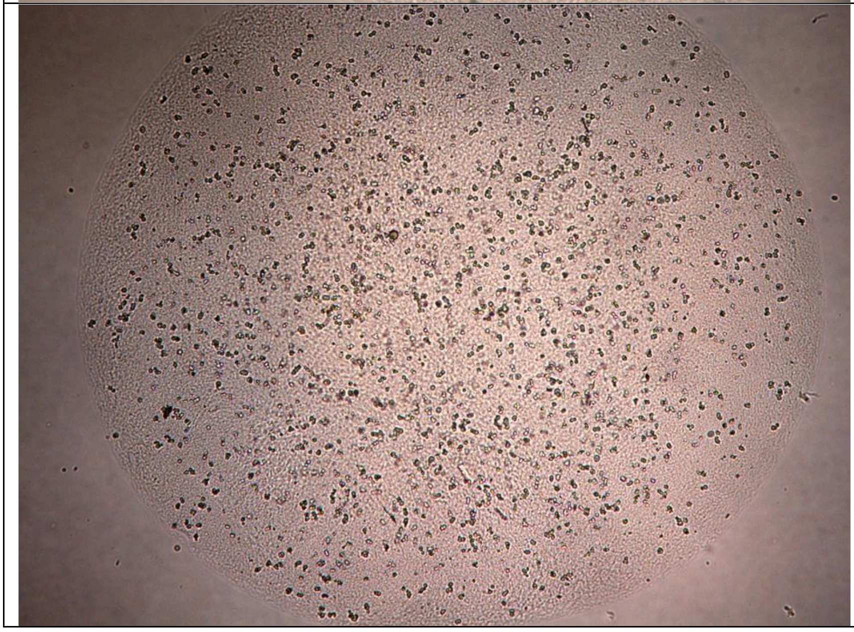 | Day 4 post printing<br><br>INS-1 832/3 cells<br>CellInk GelXA LAMININK 411<br>Seeding density<br>2×10^6/ml<br><br>4× magnification |

| | |
|---|---|
| 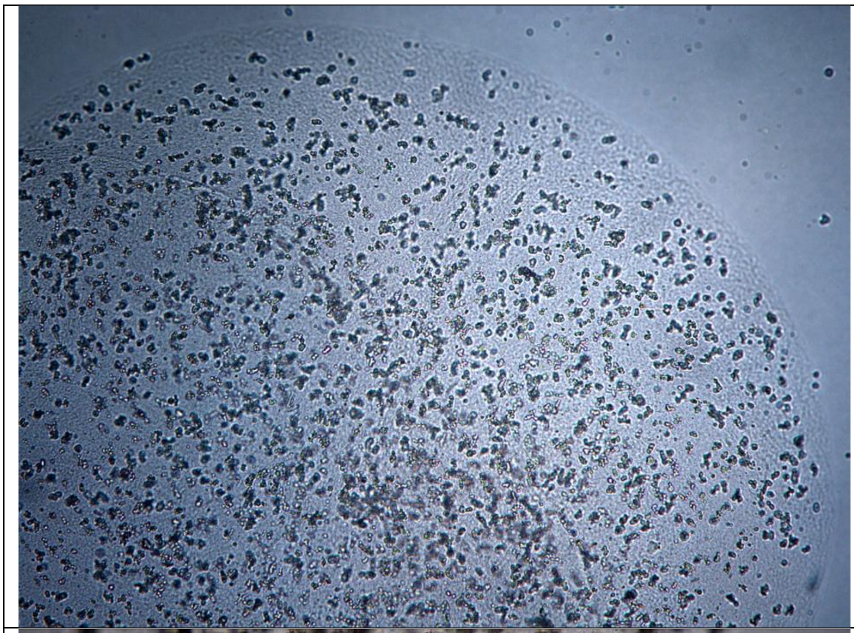 | Day 6 post printing<br><br>INS-1 832/3 cells<br>CellInk GelXA LAMININK 411<br>Seeding density 2×10^6/ml<br><br>4× magnification |
| 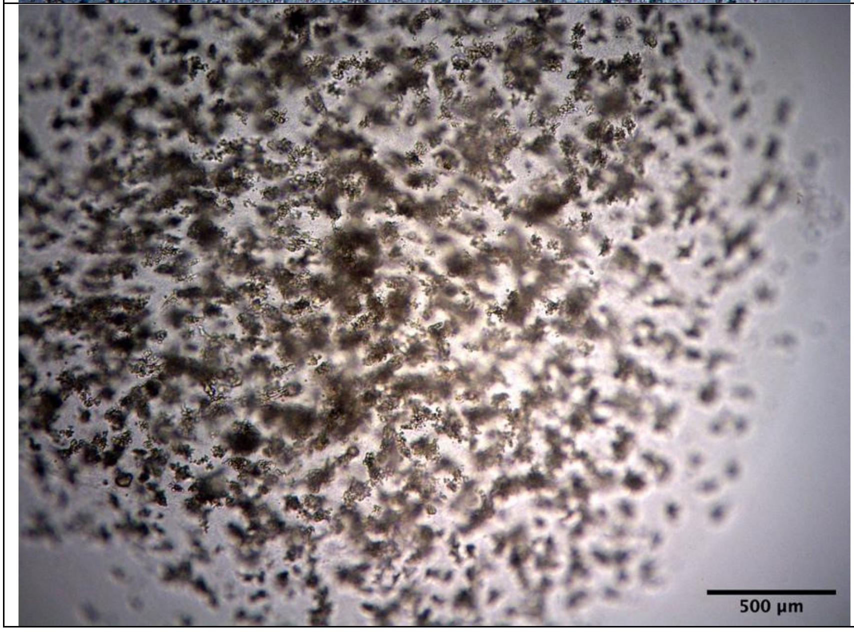 | Day 10 post printing<br><br>INS-1 832/3 cells<br>CellInk GelXA LAMININK 411<br>Seeding density 2×10^6/ml<br><br>4× magnification |

Salg, G.A. et al. Towards 3D Bioprinting of an Endocrine Pancreas: A Building-Block Concept for Bioartificial Insulin-Secreting Tissue

#### Appendix S3A
3D-Bioprinted grid structures incl. INS-1 for total RNA sequencing

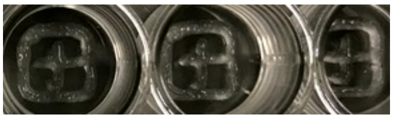

#### Appendix S3B
MA-plot RNA sequencing

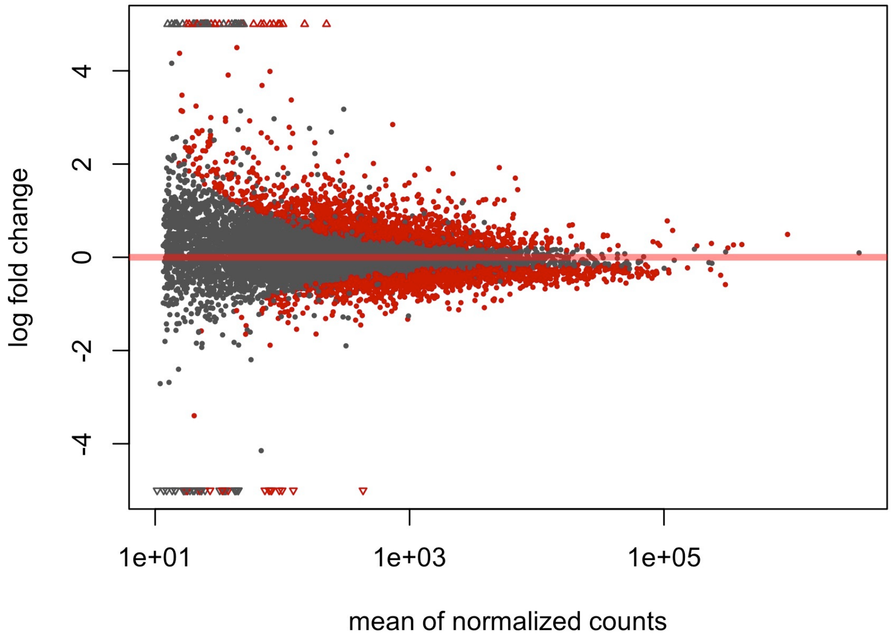

### Appendix S3C

Gene set enrichment analysis: hallmark pathways

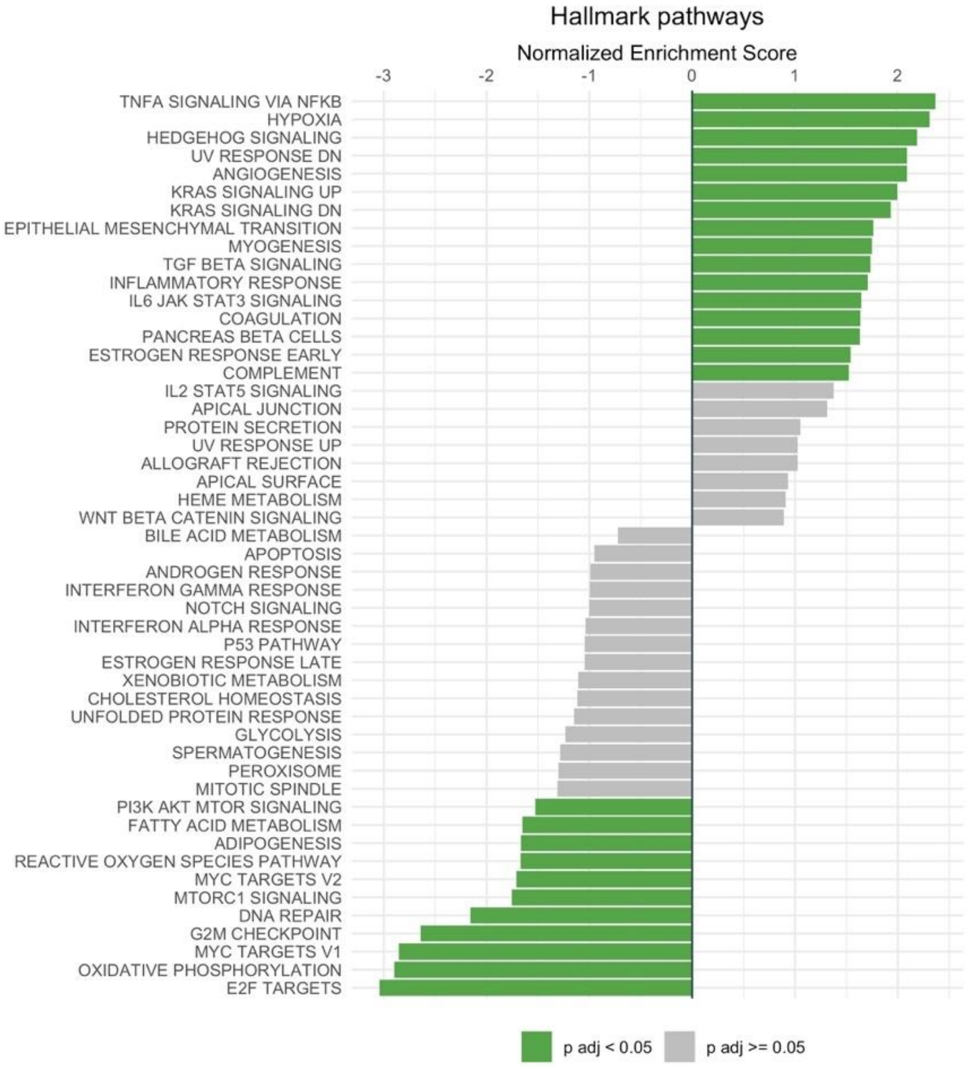

#### Appendix S3D

Ingenuity Pathway Analysis: 100 altered canonical pathways (cut-off threshold $p \leq 0.05$)

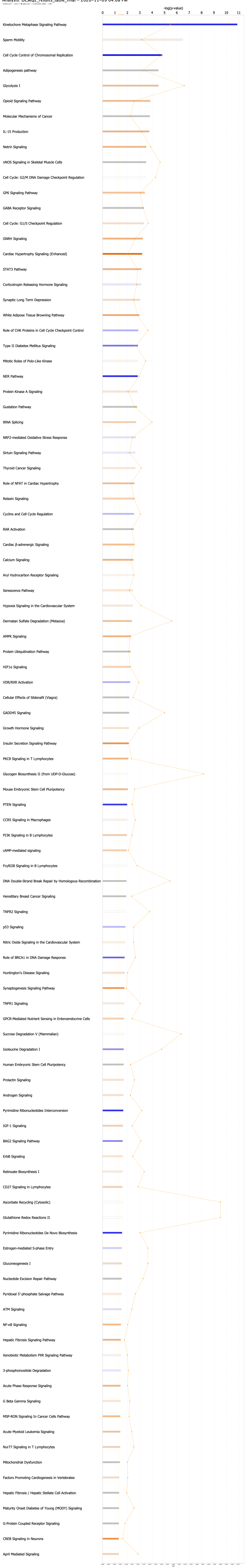


© 2000-2020 QIAGEN. All rights reserved.

#### Appendix S3E

Ingenuity Pathway Analysis: graphical summary

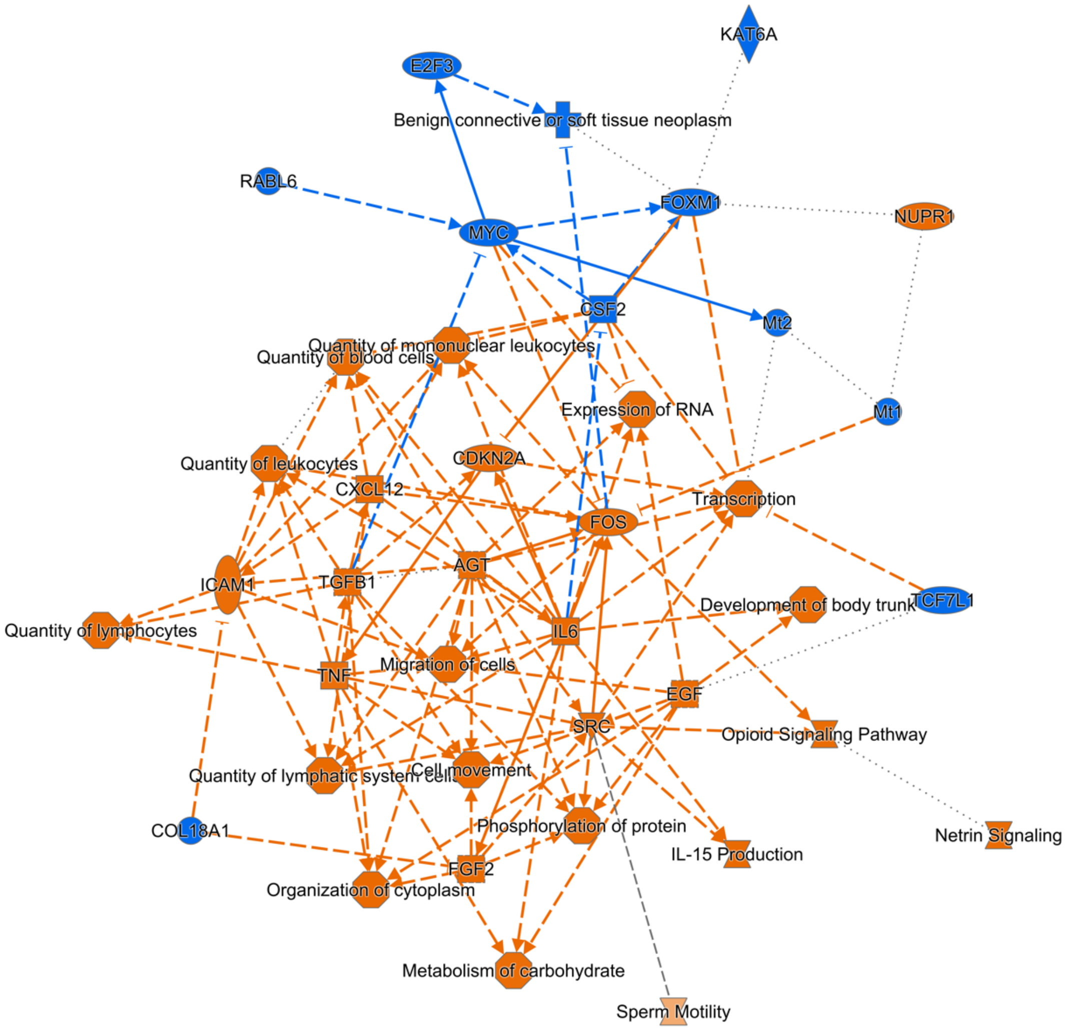

### Appendix S3F

Excerpt table: differential gene expression

| Gene_name | ID | Ensemble | baseMean | log2fold change | lfcsE | stat | p value | p adjusted |
|---|---|---|---|---|---|---|---|---|
| Insulin | Ins | ENSG00000254647 | 238211.652 | -0.129 | 0.085 | -1.522 | 0.127995498 | 0.279319244 |
| Glucagon | Gcg | ENSRNOG00000005498 | 2654.924 | 1.213 | 0.457 | 2.653 | 0.007966849 | 0.034514808 |
| Glucagon receptor | Gcgr | ENSRNOG00000036692 | 2326.087 | 1.076 | 0.101 | 10.687 | 1.17E-26 | 7.03E-24 |
| GLUT2 | Slc2a2 | ENSRNOG00000011875 | 14986.447 | 0.393 | 0.058 | 6.777 | 1.23E-11 | 7.78E-10 |
| GLUT4 | Slc2a4 | ENSRNOG00000017226 | 130.455 | 0.693 | 0.260 | 2.670 | 0.007589811 | 0.03326968 |
| SUR1, ATP binding cassette subfamily C member 8 | Abcc8 | ENSRNOG00000021130 | 2915.685 | 0.831 | 0.082 | 10.158 | 3.04E-24 | 1.42E-21 |
| MafA | Mafa | ENSRNOG00000007668 | 3771.928 | 0.158 | 0.110 | 1.433 | 0.151775038 | 0.314360904 |
| Pancreatic and duodenal homeobox 1 | Pdx1 | ENSRNOG00000046458 | 3752.708 | -0.233 | 0.090 | -2.604 | 0.009223669 | 0.038710575 |
| Neurogenic differentiation 1 | Neurod1 | ENSRNOG00000005609 | 1750.953 | 0.043 | 0.118 | 0.365 | 0.715066016 | 0.842432143 |
| Insulin like growth factor 1 receptor | Igf1r | ENSRNOG00000014187 | 361.518 | 0.638 | 0.209 | 3.052 | 0.002272084 | 0.01276656 |
| Insulin like growth factor binding protein 4 | Igfbp4 | ENSRNOG00000010635 | 529.906 | 0.750 | 0.167 | 4.491 | 7.10E-06 | 0.000100092 |
| Glycogen synthase kinase 3 beta | Gsk3b | ENSRNOG00000002833 | 678.621 | 0.555 | 0.158 | 3.517 | 0.000436008 | 0.003290919 |
| Forkhead box O1 | Foxo1 | ENSRNOG00000013397 | 353.278 | 0.642 | 0.174 | 3.685 | 0.000228759 | 0.001880739 |
| Islet amyloid polypeptide | Iapp | ENSRNOG00000012417 | 317725.627 | 0.200 | 0.059 | 3.401 | 0.000671365 | 0.004683593 |
| Insulin receptor substrate 2 | Irs2 | ENSRNOG00000023509 | 1264.762 | 0.537 | 0.106 | 5.087 | 3.64E-07 | 7.52E-06 |
| Activation transcription factor 3 | Atf3 | ENSRNOG00000003745 | 51.599 | 1.388 | 0.441 | 3.145 | 0.001658037 | 0.00991022 |
| Pyruvate dehydrogenase kinase 1 | Pdk1 | ENSRNOG00000001517 | 569.605 | 0.613 | 0.175 | 3.498 | 0.000468251 | 0.003490444 |
| Transforming growth factor beta 2 | Tgfb2 | ENSRNOG00000002418 | 92.788 | 5.784 | 0.673 | 8.592 | 8.57E-18 | 1.83E-15 |
| 6-Phosphofructo-2-kinase/fructose-2,6-bisphosphatase 3 | Pfkfb3 | ENSRNOG00000018911 | 1418.659 | 1.888 | 0.102 | 18.464 | 3.99E-76 | 2.52E-72 |
| Hypoxia inducible factor 1 subunit alpha | Hif1a | ENSRNOG00000008292 | 363.999 | 0.008 | 0.171 | 0.045 | 0.964023534 | 0.981988388 |
| Glucosidase alpha | Gaa | ENSRNOG00000047656 | 6016.747 | 0.680 | 0.080 | 8.539 | 1.36E-17 | 2.81E-15 |
| Aldolase | Aldoa | ENSRNOG00000052802 | 19051.944 | 0.696 | 0.079 | 8.863 | 7.78E-19 | 1.82E-16 |
| Oxytocin receptor | Oxtr | ENSRNOG00000005806 | 59.153 | 1.815 | 0.429 | 4.228 | 2.36E-05 | 0.000276982 |
| X-linked inhibitor of apoptosis | Xiap | ENSRNOG00000006967 | 245.367 | 0.585 | 0.266 | 2.197 | 0.028011004 | 0.092806214 |
| Caspase 3 | Casp3 | ENSRNOG00000010475 | 972.584 | -0.317 | 0.115 | -2.744 | 0.006061081 | 0.027927342 |
| Bcl2 associated X | Bax | ENSRNOG00000020876 | 3520.619 | -0.389 | 0.090 | -4.319 | 1.57E-05 | 0.000195986 |
| Bcl2 associated agonist of cell death | Bad | ENSRNOG00000021147 | 1814.186 | -0.370 | 0.093 | -3.984 | 6.77E-05 | 0.000672342 |
| Fibronectin | Fn1 | ENSRNOG00000014288 | 6783.646 | 1.698 | 0.083 | 20.548 | 7.94E-94 | 1.00E-89 |
| Ecadh | Cdh1 | ENSRNOG00000020151 | 2776.210 | 0.196 | 0.078 | 2.495 | 0.01260122 | 0.04921925 |
| VEGFA | Vegfa | ENSRNOG00000019598 | 5049.442 | 0.598 | 0.102 | 5.887 | 3.94E-09 | 1.32E-07 |
| VEGFB | Vegfb | ENSRNOG00000021156 | 863.567 | 0.502 | 0.119 | 4.215 | 2.50E-05 | 0.000290696 |
| Laminin subunit beta 3 | Lamb3 | ENSRNOG00000006025 | 479.110 | 0.721 | 0.164 | 4.387 | 1.15E-05 | 0.000150921 |
| Laminin subunit alpha 5 | Lama5 | ENSRNOG00000053691 | 181.864 | 0.672 | 0.231 | 2.902 | 0.003713368 | 0.018843066 |
| Basal cell adhesion molecule | Bcam | ENSRNOG00000029399 | 1442.506 | 0.725 | 0.137 | 5.295 | 1.19E-07 | 2.76E-06 |
| Fibroblast growth factor receptor 1 | Fgfr1 | ENSRNOG00000016050 | 529.626 | 0.953 | 0.180 | 5.307 | 1.11E-07 | 2.61E-06 |
| Fibroblast growth factor receptor 4 | Fgfr4 | ENSRNOG00000016763 | 528.082 | 0.610 | 0.146 | 4.179 | 2.92E-05 | 0.000333051 |
| Fibroblast growth factor 13 | Fgf13 | ENSRNOG00000042753 | 129.914 | 1.201 | 0.335 | 3.582 | 0.000341432 | 0.002653246 |
| Heparin binding EGF like growth factor | Hbegf | ENSRNOG00000018646 | 341.644 | 1.130 | 0.169 | 6.694 | 2.17E-11 | 1.27E-09 |
| Wnt family member 4 | Wnt4 | ENSRNOG00000013166 | 1267.758 | 0.746 | 0.099 | 7.511 | 5.88E-14 | 6.35E-12 |
| Claudin 15 | Cldn15 | ENSRNOG00000001419 | 167.689 | 1.429 | 0.248 | 5.756 | 8.59E-09 | 2.70E-07 |
| VGF nerve factor inducible | Vgf | ENSRNOG00000001416 | 5882.449 | 0.557 | 0.111 | 5.021 | 5.13E-07 | 1.02E-05 |
| Neuronal growth regulator 1 | Negr1 | ENSRNOG00000021410 | 186.823 | 0.942 | 0.226 | 4.159 | 3.19E-05 | 0.000357297 |

Salg, G.A. et al. Towards 3D Bioprinting of an Endocrine Pancreas: A Building-Block Concept for Bioartificial Insulin-Secreting Tissue

### Appendix S3G

Ingenuity Pathway Analysis: upstream regulator analysis
Glucose as upstream regulator, data set genes, effect on target molecules, and overlay of canonical pathways (insulin secretion signaling, insulin receptor signaling, glycolysis I, HIF1a signaling)

Path Designer D-glucose 1

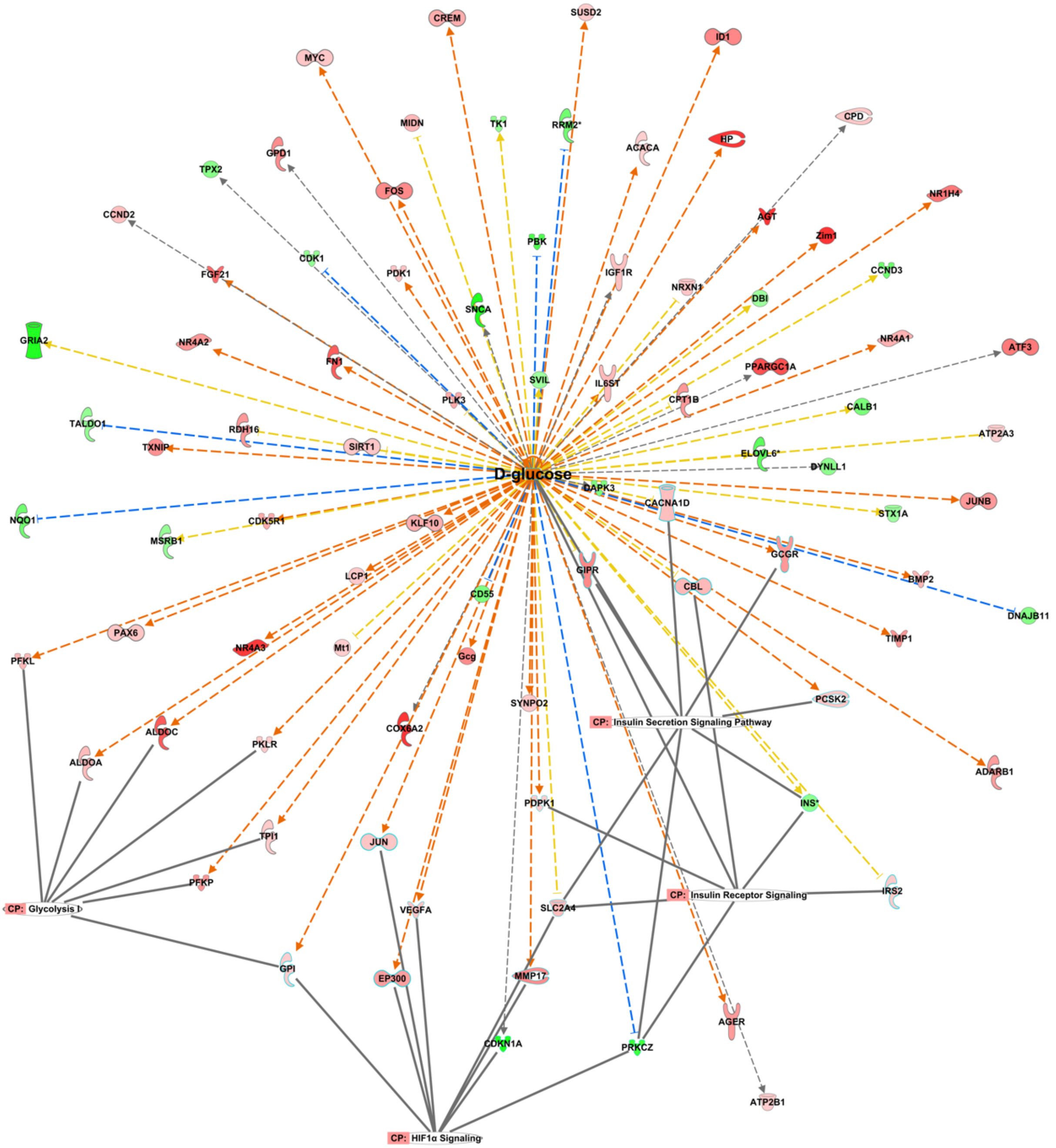

#### Appendix S4A

Vascular ingrowth and neoangiogenesis in scaffold structures
Machine learning-based vascular network analysis: Explant of PCL scaffolds from CAM assay

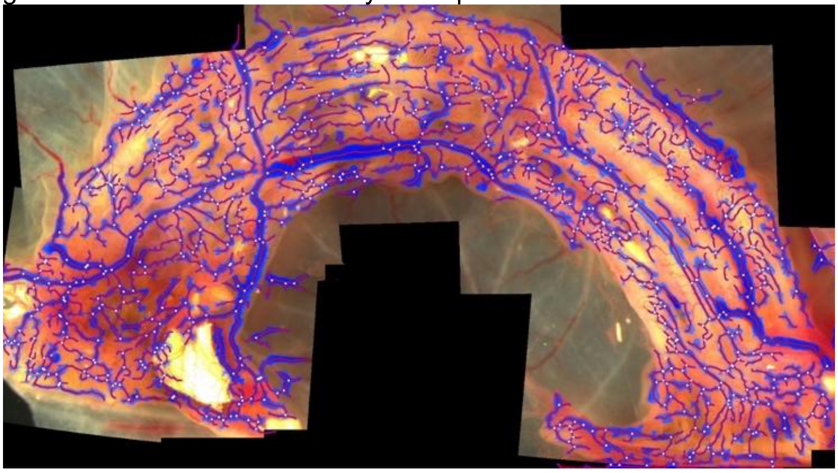

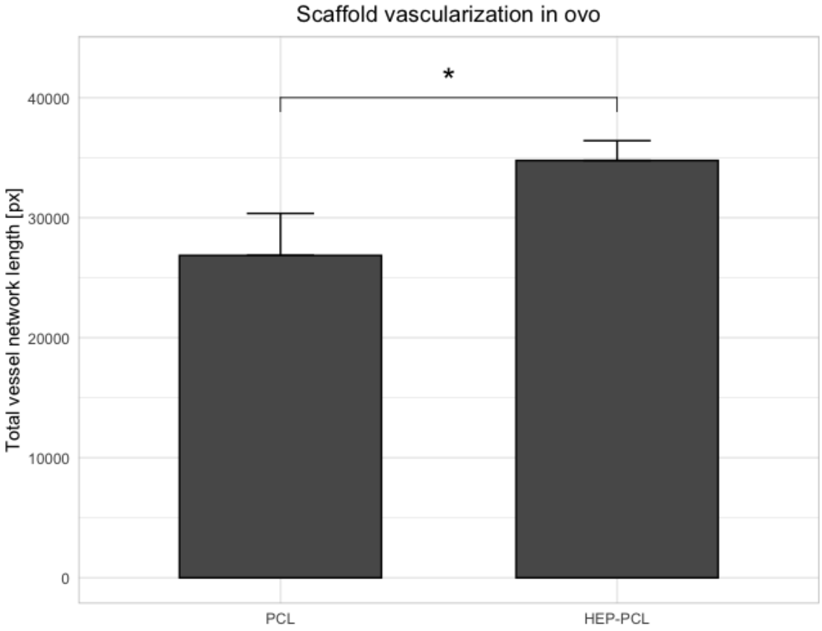

#### Appendix S4B

Chorioallantoic membrane assay: ex ovo trials, timeline

| | |
|---|---|
| 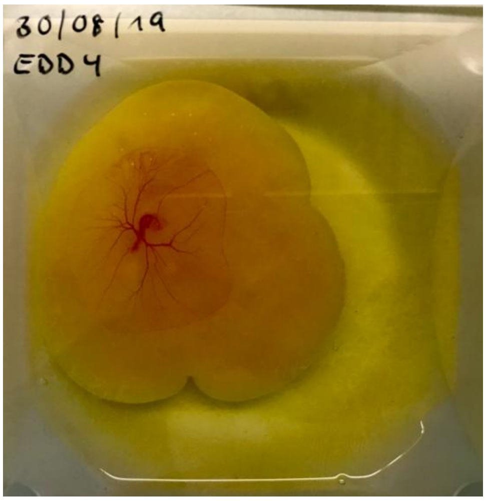  | **Embryonic development day (EDD) 4/18**<br><br>After 4 days of incubation, transfer of the viable chick embryo into culture device with glass top for observation |
| 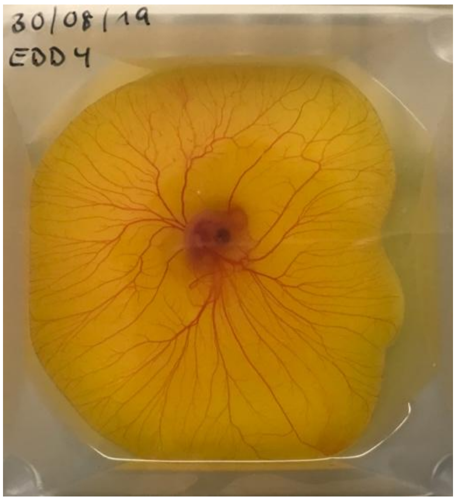  | **EDD 7/18**<br><br>Viable embryo |
| 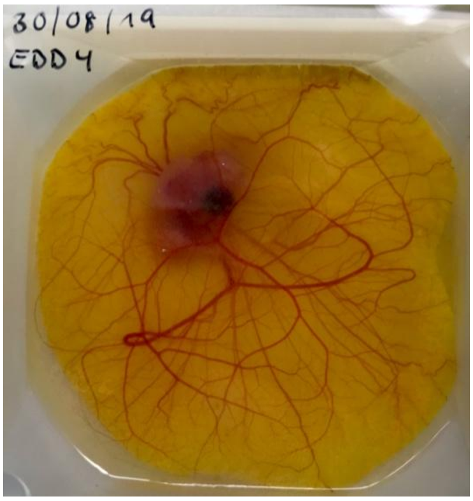  | **EDD 9/18**<br><br>Viable embryo, before implantation of solid polymer component scaffold |
| | **EDD 9/18** |

| | |
|---|---|
| 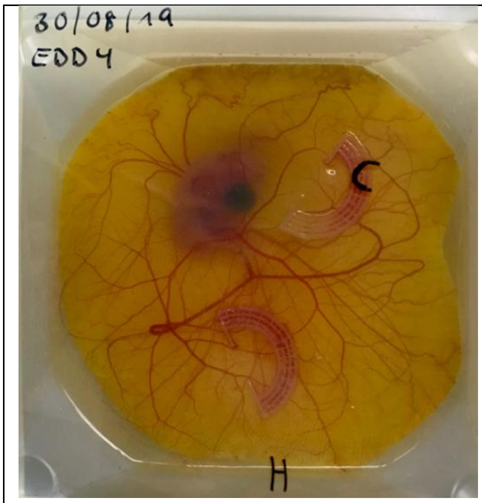 | Viable embryo, post implantation of solid polymer component scaffolds<br><br>For direct comparison, 3D-printed, sterilized untreated PCL scaffold and heparinized PCL were implanted |
| 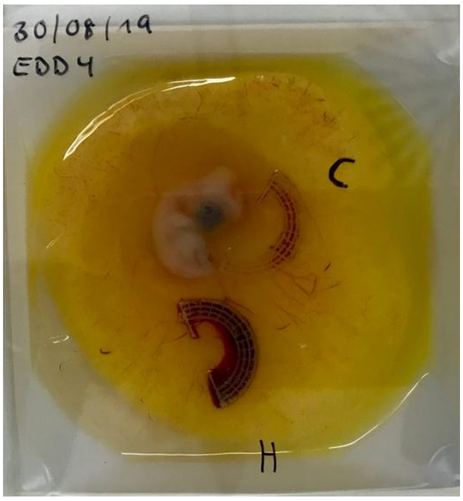 | **EDD 11/18**<br><br>Dead embryo: extensive bleeding around heparinized scaffold |

#### Appendix S4C

Immunohistochemical staining of CAM assay explant: gelatin methacrylate blend / INS-1 cells
Paraffin embedded tissue, slice thickness 5 µm, avian antiCD34 staining: bottom, periphery of xenograft; top, center of xenograft

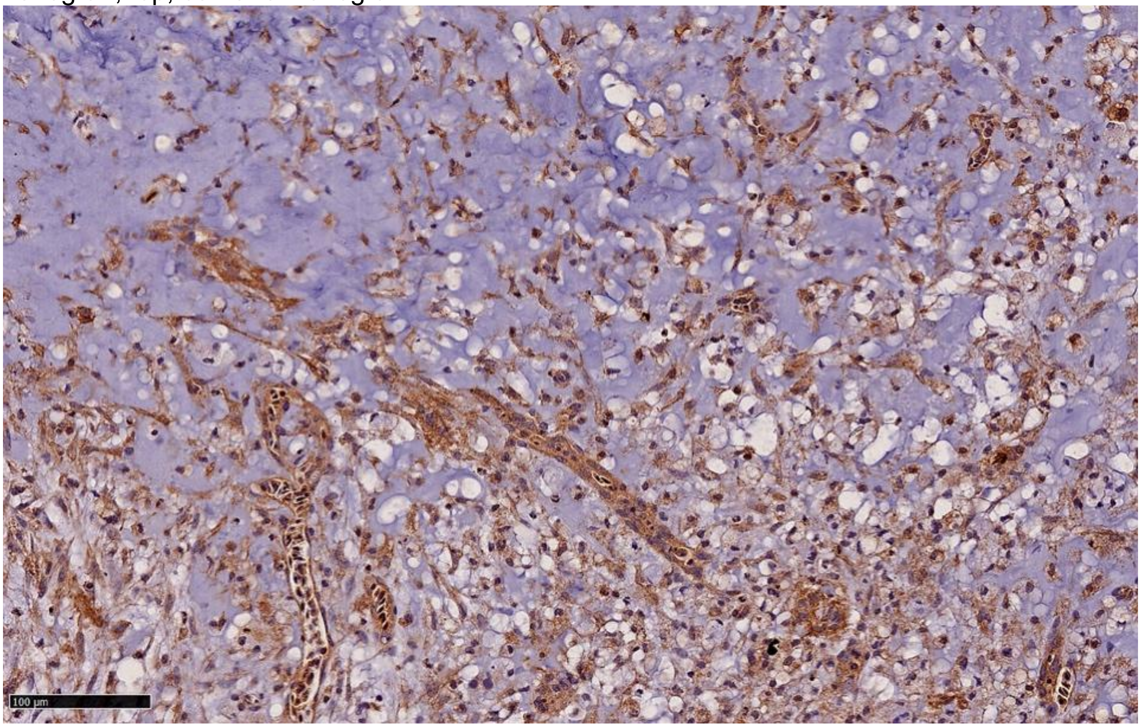

#### Appendix S5

Pixel classification with ilastik: exemplary data on spatial distribution of pseudoislets of cross-sections from bioprinted xenografts explanted from CAM (anti-insulin immunohistochemistry)

Pseudoislet segmentation (Insulin$^+$ staining) using ilastik pixel classification with overlay of hydrogel graft area (1.3 mm$^2$, 1.8% pseudoislet area). Cross section at 20 µm (from graft base).

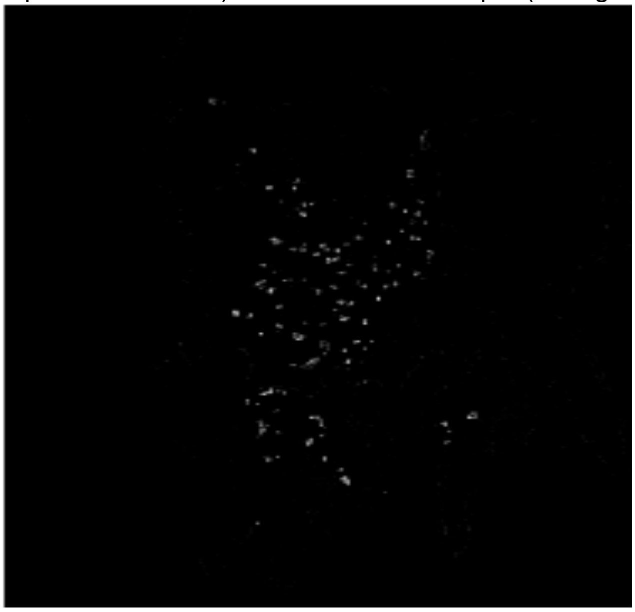

Pseudoislet segmentation (Insulin$^+$ staining) using ilastik pixel classification with overlay of hydrogel graft area (1.8 mm$^2$, 2.1% pseudoislet area). Cross section at 40 µm (from graft base).

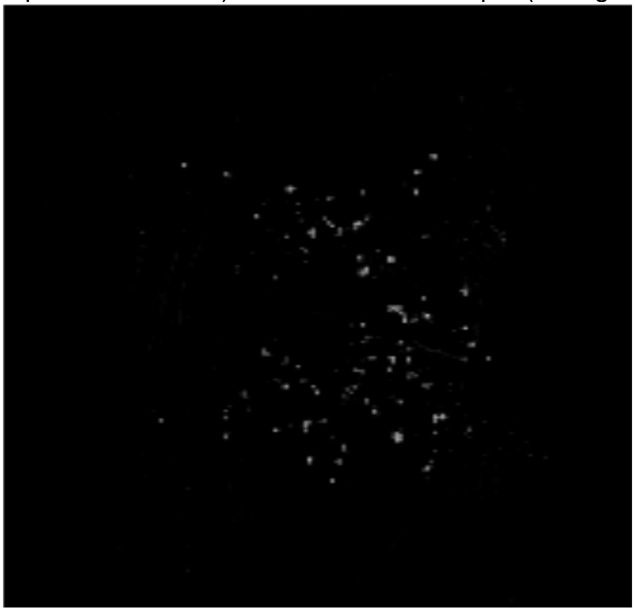

Pseudoislet segmentation (Insulin$^+$ staining) using ilastik pixel classification with overlay of hydrogel graft area (6.5 mm$^2$, 2.1% pseudoislet area). Cross section at 60 µm (from graft base).

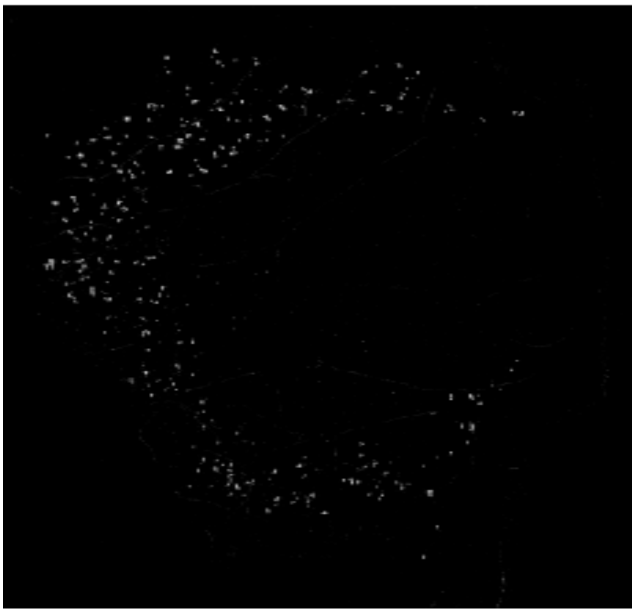

Pseudoislet segmentation (Insulin+ staining) using ilastik pixel classification with overlay of hydrogel graft area (10 $mm^2$, 0.1% pseudoislet area). Cross section from xenograft with 15 s UV crosslinking (405 nm) after bioprinting

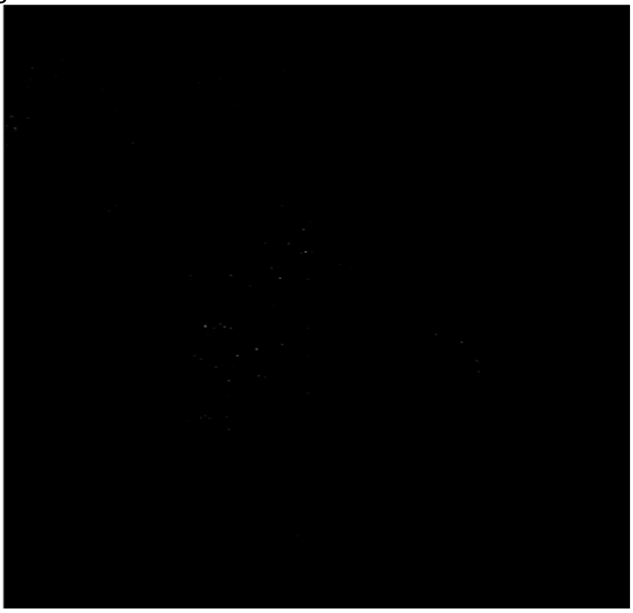

Pseudoislet segmentation (Insulin+ staining) using ilastik pixel classification with overlay of hydrogel graft area (21.6 $mm^2$, 0.6% pseudoislet area). Cross section from large xenograft with 15 s UV crosslinking (405 nm) after bioprinting

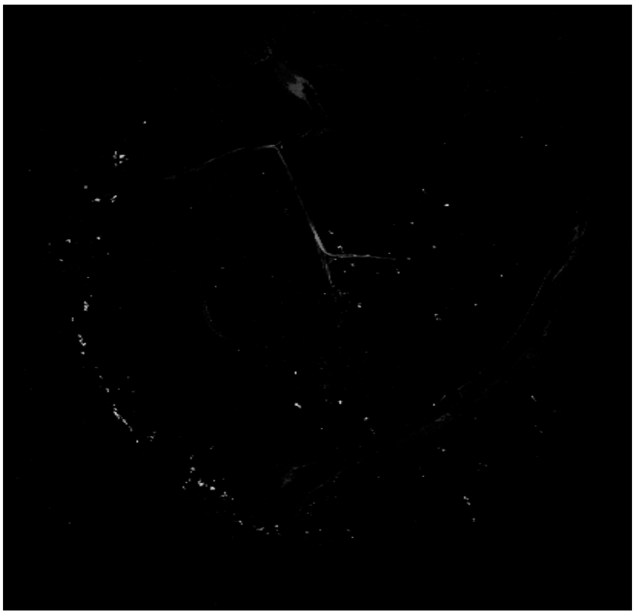

#### Appendix S6

GSIS: experimental workflow

INS-1 cells were either growing in monolayer (2D samples) or embedded in bioprinted LAMININK 411 hydrogels (3D samples). To investigate the influence of endothelial cells on INS-1 cells, hydrogels containing a 1:2 co-culture of INS-1 and HUVEC were printed. GSIS was performed and cells were counted using fluorescence microscopy.

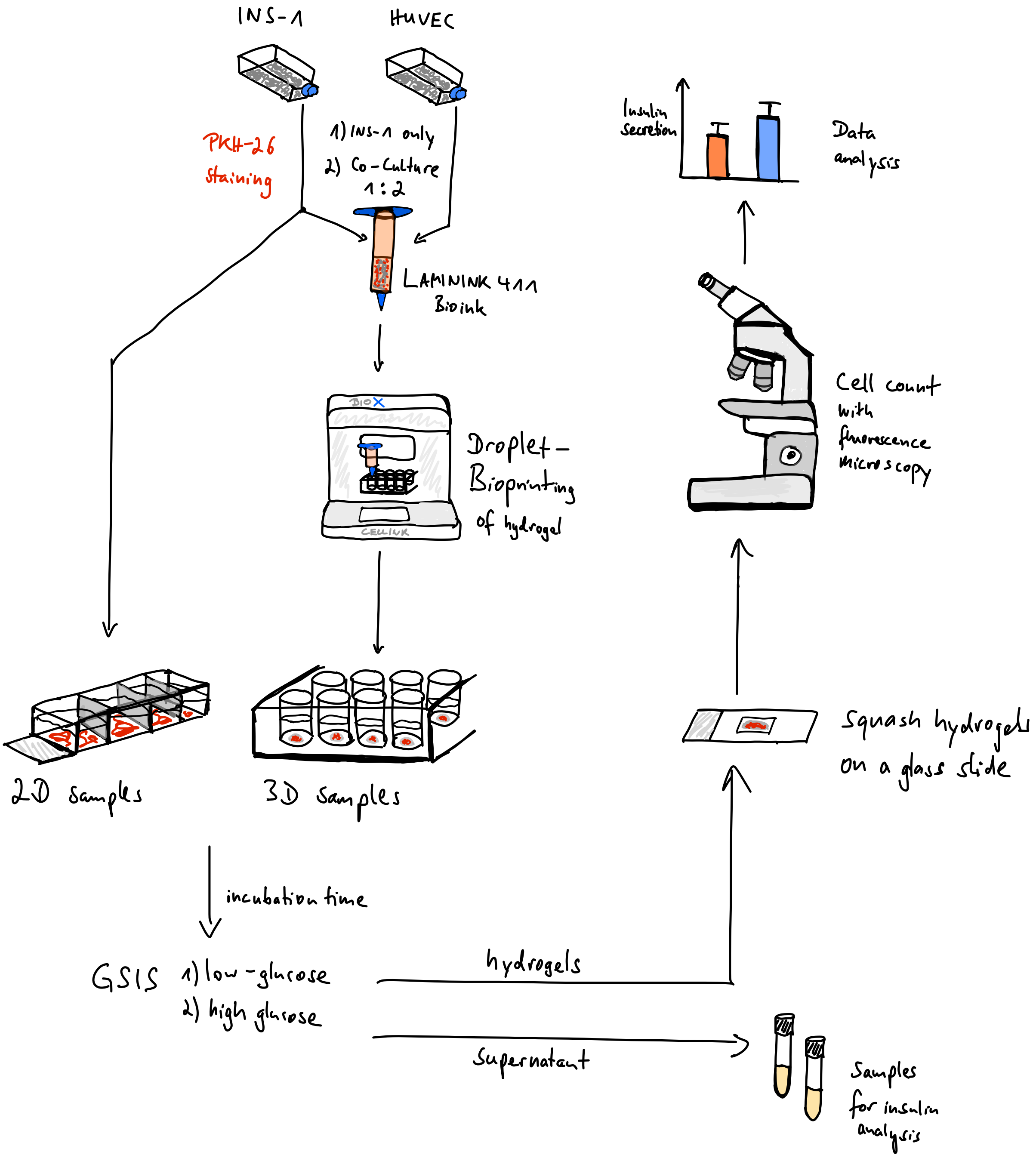

Salg, G.A. et al. Towards 3D Bioprinting of an Endocrine Pancreas: A Building-Block Concept for Bioartificial Insulin-Secreting Tissue

GSIS: culture conditions for INS-1 cells, 2D monolayer vs. 3D hydrogel

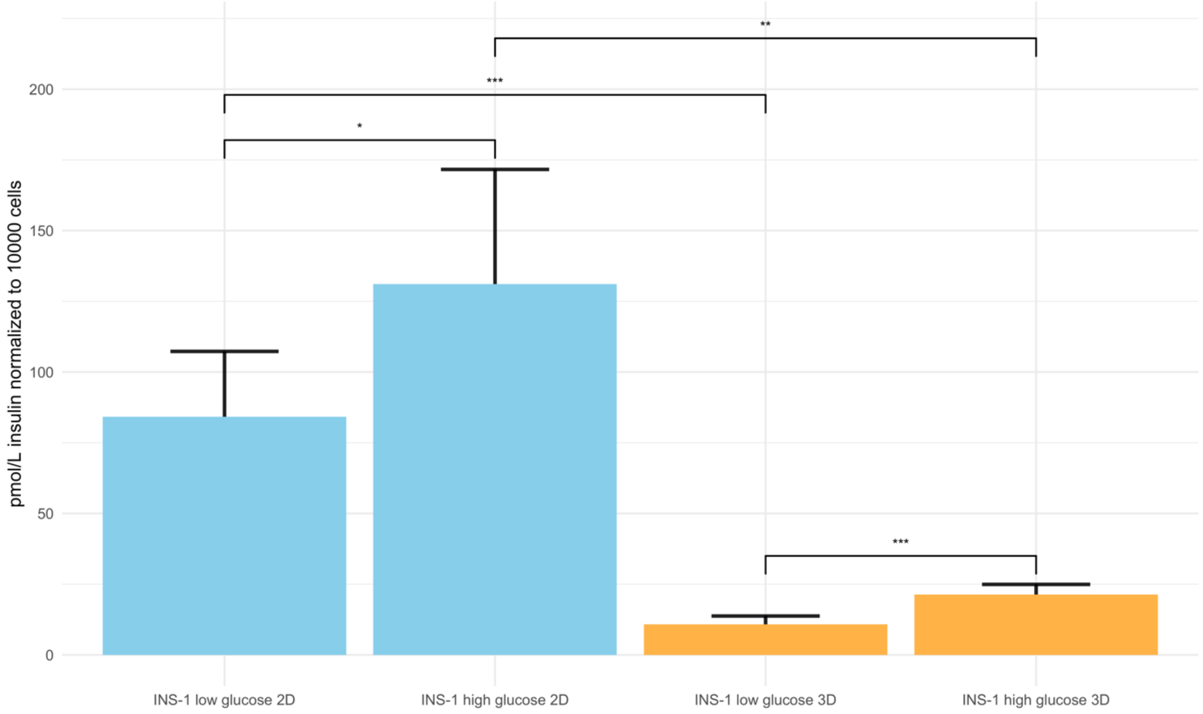

### Appendix S7

In silico analysis: parameters for computer-aided structure screening

| Parameter | Value |
|---|---|
| ***Oxygen*** | |
| Initial and inflow concentration | 0.1305 mol/m³ (equ. 90 mmHg)<br>0.232 mol/m³ (equ. 160 mmHg)<br>0.3915 mol/m³ (equ. 270 mmHg) |
| Diffusion through aqueous media | $3.0 \times 10^{-9}$ m²/s * |
| Diffusion through hydrogel | $2.5 \times 10^{-9}$ m²/s * |
| Diffusion through Langerhans islet | $2.0 \times 10^{-9}$ m²/s * |
| ***Glucose*** | |
| Initial and inflow concentration | 5 mol/m³<br>10 mol/m³<br>15 mol/m³<br>25 mol/m³ |
| Diffusion through aqueous media | $9.0 \times 10^{-10}$ m²/s * |
| Diffusion through hydrogel | $6.0 \times 10^{-10}$ m²/s * |
| Diffusion through Langerhans islet | $3.0 \times 10^{-10}$ m²/s * |
| ***Insulin*** | |
| Initial and inflow concentration | 0 mol/m³ |
| Diffusion through aqueous media | $1.5 \times 10^{-10}$ m²/s * |
| Diffusion through hydrogel | $1.0 \times 10^{-10}$ m²/s * |
| Diffusion through Langerhans islet | $0.5 \times 10^{-10}$ m²/s * |
| ***Islet of Langerhans*** | |
| Radius | 50 µm<br>75 µm<br>150 µm<br>250 µm |
| ***Hydrogel*** | |
| Shell thickness | 0 µm<br>50 µm<br>100 µm<br>300 µm<br>500 µm<br>600 µm<br>700 µm<br>800 µm<br>1000 µm |

*As described by Buchwald. et al. (2011)
